# Expectations affect the perception of material properties

**DOI:** 10.1101/744458

**Authors:** Lorilei M. Alley, Alexandra C. Schmid, Katja Doerschner

**Affiliations:** Justus Liebig University Giessen; Bilkent University

## Abstract

Many objects that we encounter have ‘typical’ material qualities: spoons are hard, pillows are soft, and jell-O dessert is wobbly. Over a lifetime of experiences, strong associations between an object and its typical material properties may be formed, and these associations not only include how glossy, rough or pink an object is, but also how it behaves under force: we expect knocked over vases to shatter, popped bike tires to deflate, and gooey grilled cheese to hang between two slices of bread when pulled apart. Here we ask how such rich visual priors affect the visual perception of material qualities and present a particularly striking example of expectation violation. In a cue conflict design, we pair computer-rendered familiar objects with surprising material behaviors (a linen curtain shattering, a porcelain teacup wrinkling, etc.) and find that material qualities are not solely estimated from the object’s kinematics (i.e. its physical (atypical) motion while shattering, wrinkling, wobbling etc.); rather, material appearance is sometimes “pulled” towards the “native” motion, shape, and optical properties that are associated with this object. Our results, in addition to patterns we find in response time data, suggest that visual priors about materials can set up high-level expectations about complex future states of an object and show how these priors modulate material appearance.

## Introduction

The material an object is made of endows it with a certain utility or function: chairs are usually rigid, in order to afford sitting, spoons are hard to afford eating, and towels are soft and absorbent to afford drying. The material choices for many types of man-made objects (keys, cups, cushions) tend to be restricted, and thus many objects that we encounter have a ‘typical’ material: bookshelves are made from wood, pillows from cloth and down, and keys from metal. Over a lifetime of interacting with objects, we learn not only what they look like, but also how their material ‘behaves’ under different types of forces. For example, we know that porcelain vases shatter when knocked over, that soft jell-O wobbles when poked, and that rubber balls bounce when thrown at the wall. Strong associations are formed between the object’s shape and its optical and mechanical material properties.

Observers rely on these associations when estimating material qualities, which can lead to paradoxical findings. For example, physically, the softness of an object (compliance) is independent of its optical properties (color, gloss or translucency), and with sophisticated production techniques, it is possible to manufacture soft objects with any set of optical properties. Yet, when judging the softness of an object in static images, research has shown that that optical characteristics of the material affected observers’ judgements, e.g. a velvety cube was perceived as softer than a chrome cube of the same shape and size (Paulun, Schmidt, van Assen, & Fleming, 2017; Schmidt, Paulun, van Assen & Fleming, 2017). To account for these results, Paulun et al., (2017) propose an indirect route to perception, where image cues activate the memory of a particular material along with its learned associations, including its typical mechanical properties (e.g. ‘this looks like honey - so it’s probably quite runny.’). They suggest that this route operates in parallel with a more direct one, where image cues directly convey something about the properties of the material (e.g. Paulun et al., 2017, Paulun, Kawabe, Nishida, & Fleming, 2015, van Assen, Wijntjes, & Pont, 2016, van Assen, Barla, & Fleming, 2018, Marlow & Anderson, 2016, see Figure 1 for an illustration).

**Figure 1.**
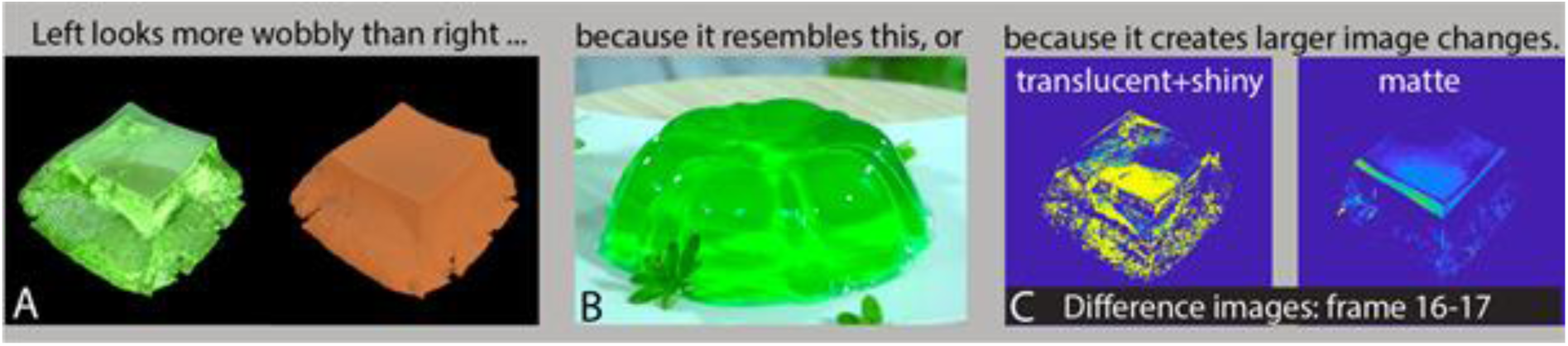
Contribution of prior associations and image cues on perceived material qualities. The role of predictions or associative mechanisms in material perception is not well understood. A. The perception of material qualities (such as gelatinousness) can be influenced by prior associations between dynamic optics, shape, and motion properties. **A.** Watching the green (left) object deform may evoke an association with green Jell-O (**B**.), and may therefore be perceived as wobblier and more gelatinous than the matte object, despite both objects wobbling in identical ways (as shown in Supplementary Movies S1 and S2). **C.** Alternatively, the green object may be perceived as wobblier due to larger image differences between frames, and potentially higher motion energy, as illustrated on the right (Doerschner et al., 2011, Doerschner, Kersten, & Schrater, 2011)). A combination of associative and modulatory mechanisms is also possible. The difference in motion energy in images of the translucent object in panel C. is about seven (6.8) times larger than that of the matte one, purely due to the difference in optical properties between these two objects.

Learned associations about material properties are not only evoked by the optical characteristics of an object but may also be elicited by its shape: Experience with soft materials or liquids appears to create strong associations between shape deformations and perceived material qualities (e.g. Paulun et al., 2017, Schmidt et al., 2017, Paulun et al., 2015, van Assen et al., 2016, van Assen et al., 2018, Schmid & Doerschner, 2018, Kawabe, 2018, Mao, Lagunas, Masia, & Gutierrez, 2019, Bi, Jin, Nienborg, & Xiao, 2019, Kawabe, Maruya, Fleming, & Nishida, 2015). Similarly, if a shape reminds the observer of a specific object, its typical mechanical material properties might also become activated. Consequently, recognizing an object should cause observers to activate strong predictions about the object’s material properties, and how it should behave under physical forces. Here we ask whether these predictions in turn affect how we perceive material properties?

The human brain uses prior knowledge to continuously generate predictions about visual input in order to make quick decisions and guide our actions (see Kveraga, Ghuman, & Bar, 2007 for a review). Predictions about material properties should be no exception to this: To avoid small daily disasters, we need to be able to predict how bendable and heavy a cup is before picking it up. How prior knowledge is combined with visual input has been investigated in several areas of vision. For example, researchers have studied how color memory for objects influences perceived color (e.g. Hansen, Olkkonen, Walter, & Gegenfurtner, 2006; Olkkonen & Allred, 2014), or how memory modulates the perception of motion direction or dynamics of binocular rivalry (Scocchia, Valsecchi, Gegenfurtner, & Triesch, 2013 & Scocchia, Valsecchi, & Triesch, 2014, Chang & Pearson, 2018). Within the framework of Bayesian models (e.g. Landy, Maloney, Johnston, & Young, 1995; Ernst & Banks, 2002; Hillis, Ernst, Banks, & Landy, 2002; Kersten, Mamassian, & Yuille, 2004) the influence of prior knowledge on the integration of different types of sensory input has been looked at more formally. In particular, cue conflict scenarios have proven extremely useful to generate insights about the complex interplay of prior selection and the weighting of sensory input in the perception of object properties (e.g. Knill, 2007). We will use an experimental paradigm analogous to cue conflict by juxtaposing indirect (prior) and direct (sensory) information in the perception of material properties, to formally test whether expectations about an object’s mechanical properties are generated.

Violating the expected mechanical properties of materials necessarily involves image motion. Figure 2 illustrates this. Shown below are three frames from an animation by Florent Porta. The first panel sets up the viewer’s expectations about the objects material properties, (i.e. the balloon will pop when it comes into contact with the spines of the cactus). As the movie proceeds, our expectations about the event-to-unfold are violated, and the viewer is quite surprised when the cactus pops like the balloon would. In the present study we directly test how such expectations affect the perception of material properties, by comparing material perception for falling objects that deform in surprising and expected ways.

**Figure 2.**
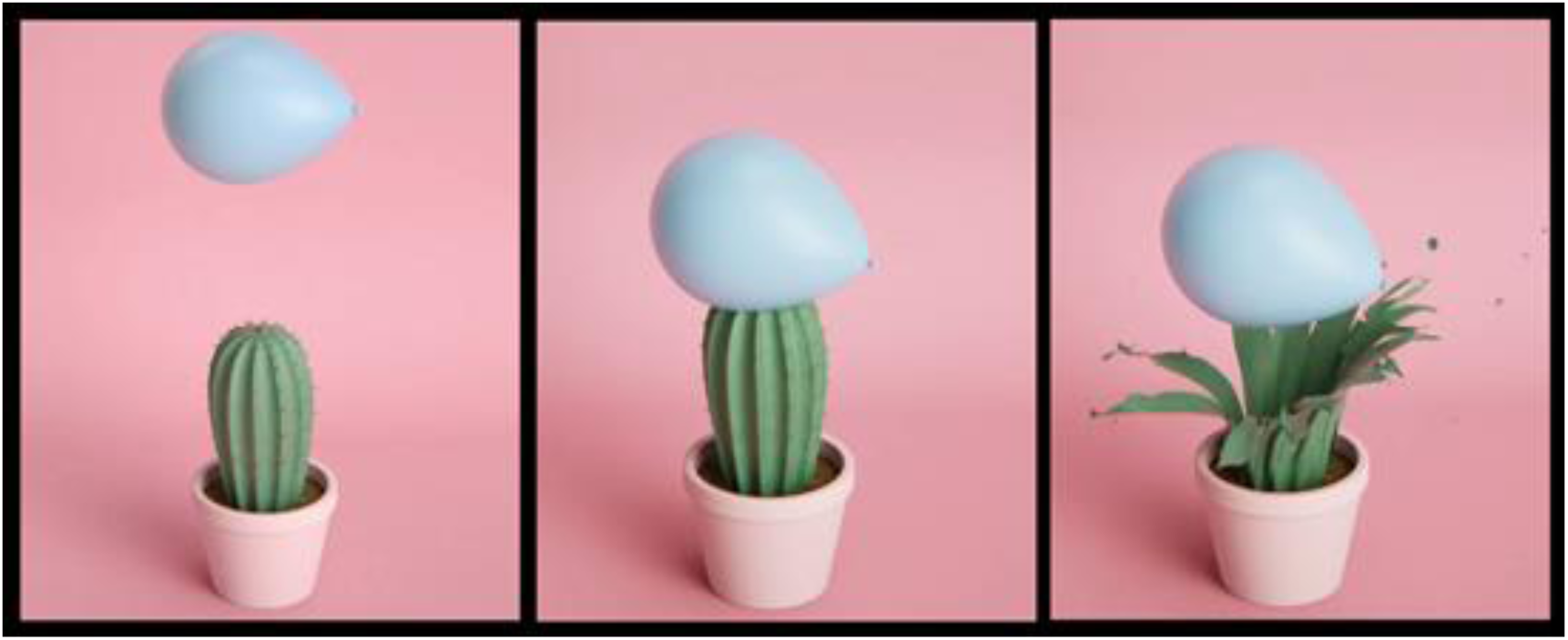
Three frames from “Preposterous” by Florent Porta. Artists have played with our expectation of how objects and their materials should behave. In this study, we compare material perception for falling objects that deform in surprising and unsurprising (i.e. expected) ways. Retrieved from https://vimeo.com/191444383.

We anticipate our results using the Bayesian framework for analogy. Suppose the task of an observer, watching the movie in Figure 2, was to rate the cactus’ rigidness. If the observer recognizes the object as a cactus, it is probably safe to assume that observers have a strong prior belief about how rigid cacti are, but it is also possible that other, weaker, prior beliefs exist. When confronted with a popping cactus, the visual system may either veto all cues that previously suggested that the cactus was rigid (e.g. Landy, Maloney, Johnston, & Young, 1995) and judge the cactus as very soft, or it may be that the visual system down-weighed cues to rigidness (Knill, 2007a). If cue vetoing occured, we would expect to find no difference in the perception of mechanical material properties (e.g. perceived rigidness in our example) between objects that deform in surprising (popping cactus) and expected (popping balloon) ways– because in either case the strongest cue is used (i.e. the visual sensory input which shows the cactus is popping and thus really not that rigid). If, however, a down-weighing of the visual input occurred, the visual prior (on the typical rigidness of cactii) would exert an influence when observers rated the rigidness of the popping cactus and we would find they are rated as somewhat more rigid than popping baloons. The Bayesian framework predicts another result:when expectations are violated, as in the exploding cactus scenario, the visual system needs to update its internal model of the world (the generative model, e.g. see Kersten et al., 2004), in order to minimize the prediction error for future tasks. This updating (or prediction error correction) is thought to be a reiterative process, which may take time to complete. In fact, a recent study by (Urgen & Boyaci, 2019) models an individuation task and shows that for surprising trials, the model predicted longer duration thresholds. By analogy, we expect to find that it takes observers longer to perform perceptual tasks when judging material attributes of surprisingly deforming objects (e.g. popping cactii) than ones that deform as expected (e.g. popping balloons).

In this study, we use a violation of expectation paradigm to investigate how predictions about object deformations based on object knowledge influence how we perceive the material of an object. In order to manipulate object familiarity, we rendered two types of objects: Familiar objects, for which there exist strong predictions about their mechanical material properties, and Novel, unfamiliar objects, for which no strong predictions should exist. For each of these object types, we rendered two motion sequences that showed how objects deformed when being dropped on the floor: an ‘Expected’ sequence, in which objects’ deformations were consistent with the observer’s prior beliefs about the mechanical properties of the material, and a ‘Surprising’ sequence in which they were not. The Novel objects inherited their optical and motion properties from a corresponding Familiar object. Participants rated the objects in these movies on various material attributes. While we expected to find differences in ratings and response times between Expected and Surprising events for Familiar objects (see above), we also expected to find differences - although attenuated - for Novel shapes. Note that we expected *some* rating differences between Expected and Surprising deformations also to occur for the unfamiliar, Novel shapes due to their optical cues alone and/or the fact that they all were bounded 3D objects, which might have elicited a prior expectation (e.g. a rigidity prior) about these objects’ mechanical material properties. We included two additional experimental conditions in which observers rated the same attributes on static images that showed each familiar and novel object during the first and last frame of the motion sequence, in order to explicitly assess for each object (familiar and novel) existing priors on material properties, as well as the influence of shape recognizability (i.e. after the object had deformed) on ratings, respectively.

## Methods

### Stimuli

#### Objects

We used two types of objects: Familiar and Novel. Figure 3 gives an overview of all objects used in the experiment. In order to create a stimulus set with broad range of typical material classes, we choose 15 Familiar objects belonging to one of 5 material mechanics: non-deforming (wooden chair, metal key, metal spoon), wobbling (red and green jell-O, custard), shattering (wineglass, terracotta pot, porcelain teacup), wrinkling (linen, velvet, and silk curtain) and splashing (milk, honey, and water drops). All objects were rendered (Blender 2.77a, Stichting Blender Foundation, Amsterdam, NL) with their typical optical material properties, e.g. metallic-looking key, green transparent jell-O, or a silky appearing curtain). Objects were located in a room that had brown walls and a hardwood, polished floor, and were illuminated by the “Campus” environment map (Debevec, 1998). The 3D meshes of the Familiar objects were obtained or adapted from TF3DM (www.tf3dm.com) and TurboSquid (www.turbosquid.com), or created by hand. Unfamiliar shapes (‘Glavens’, Phillips, 2004) inherited their optical and mechanical properties from the corresponding Familiar object.

**Figure 3.**
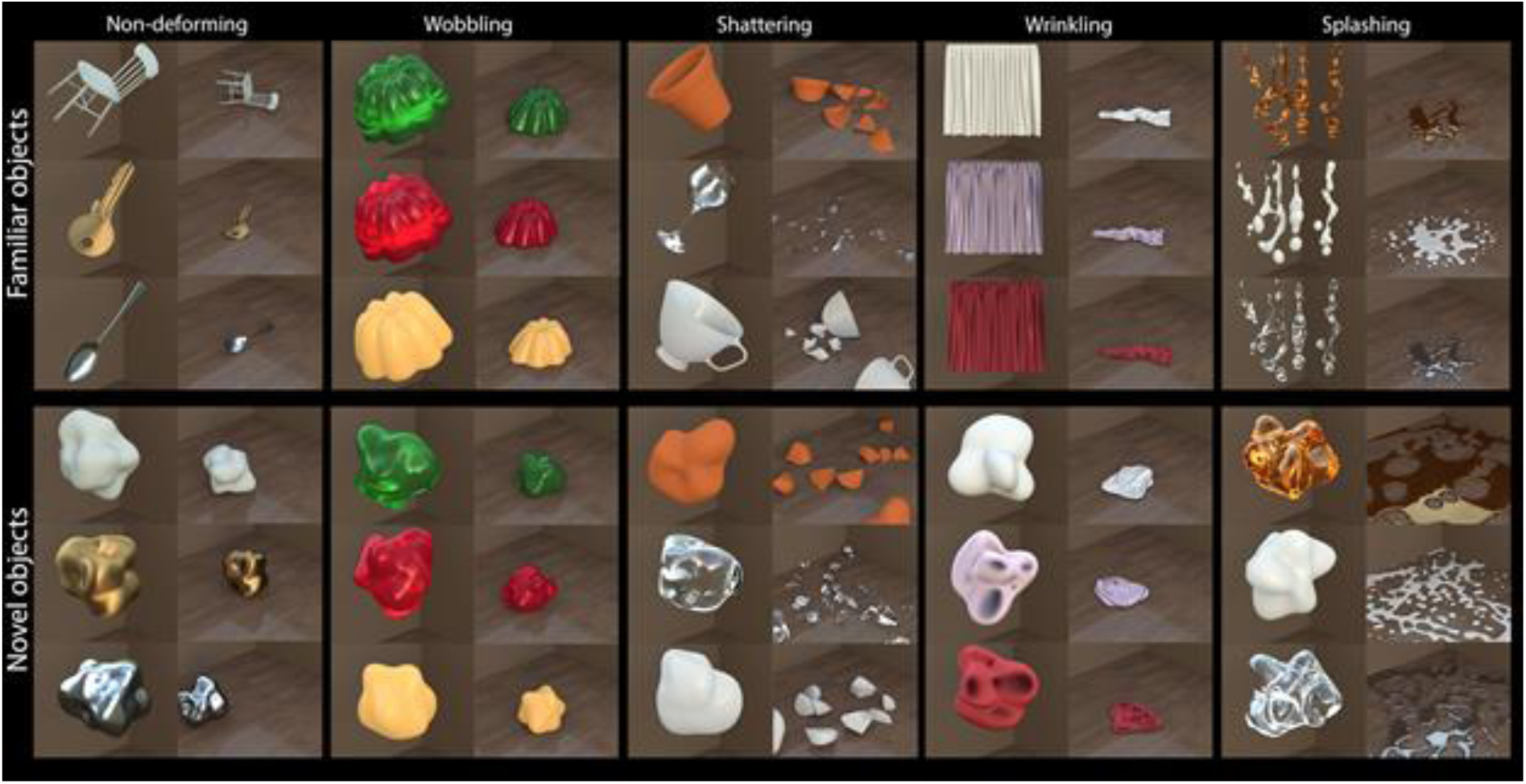
Stimuli Expected Motion. Shown are all 15 familiar (top) and corresponding novel objects (bottom) used in the experiments. Objects are organized according to their typical material kinematics, i.e. how they deform under force. Note that individual scenes are scaled to maximize the view of the object (First frame, left columns), or in order to give a good impression of the material kinematics (Last frame, right columns).

The objects were rendered at approximately the same size as each other, so that for example the key was the same “physical” (simulated) size as the chair, even though in real life chairs are larger than keys. We chose to do this so that the objects would hit the ground at the same time, and behave in a similar way under gravity. This is important for some of our analyses (see Analysis - Response time).

#### Deformations

For each object, we rendered short movies that showed an object falling from a height and interacting with the ground. In order to manipulate surprise in our experiment, an object could either behave as expected, e.g. a glass would shatter, or it could inherit the mechanical material behavior from another object, e.g. milk drops would stay rigid upon impact (Figure 4B shows two more examples of surprising material ‘behaviors’, also see Supplementary Figure 1). We created corresponding Expected and Surprising movies for Novel objects (Figure 3, Figure 4C & Supplementary Figure 1). Table 1 shows which objects inherited which material mechanics in the Surprising condition. All experimental movies can be downloaded at https://doi.org/10.5281/zenodo.2542577.

**Table 1.**
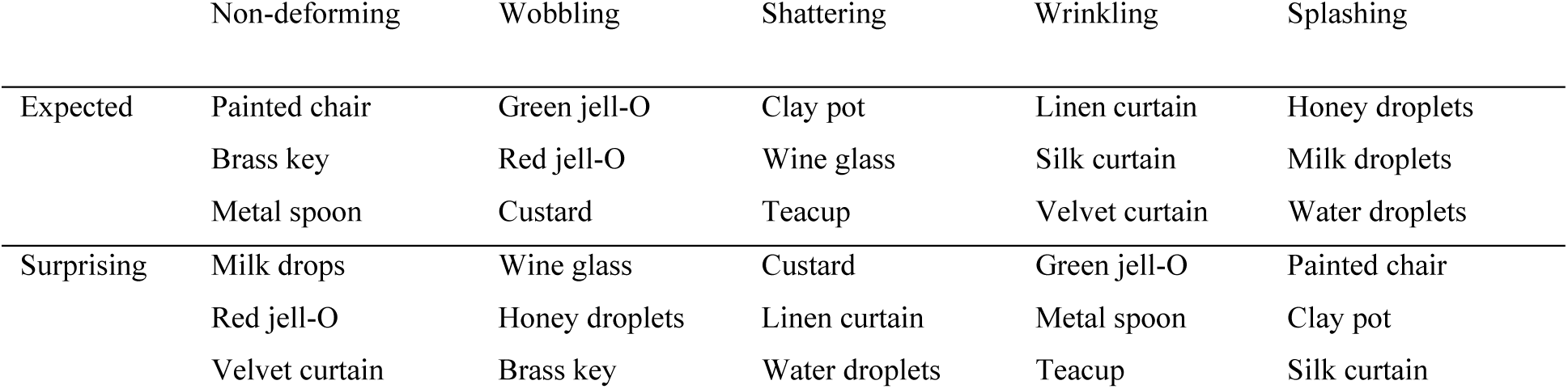
Overview of objects and conditions in the experiment. Columns show the five categories of typical material mechanics (material class) in the experiment; rows show which objects occurred in the Expected and Surprising conditions. Every object in the Expected condition would also appear exactly once in the Surprising condition, where it would be rendered with very atypical mechanical properties, e.g. the silk curtain would splash (see Figure 3 and Supplementary Figure 1 for corresponding renderings). Novel objects had no familiar shape but had the same optical and mechanical material properties as the Familiar objects in this table.

|  | Non-deforming | Wobbling | Shattering | Wrinkling | Splashing |
| --- | --- | --- | --- | --- | --- |
| Expected | Painted chair | Green jell-O | Clay pot | Linen curtain | Honey droplets |
|  | Brass key | Red jell-O | Wine glass | Silk curtain | Milk droplets |
|  | Metal spoon | Custard | Teacup | Velvet curtain | Water droplets |
| Surprising | Milk drops | Wine glass | Custard | Green jell-O | Painted chair |
|  | Red jell-O | Honey droplets | Linen curtain | Metal spoon | Clay pot |
|  | Velvet curtain | Brass key | Water droplets | Teacup | Silk curtain |

**Figure 4.**
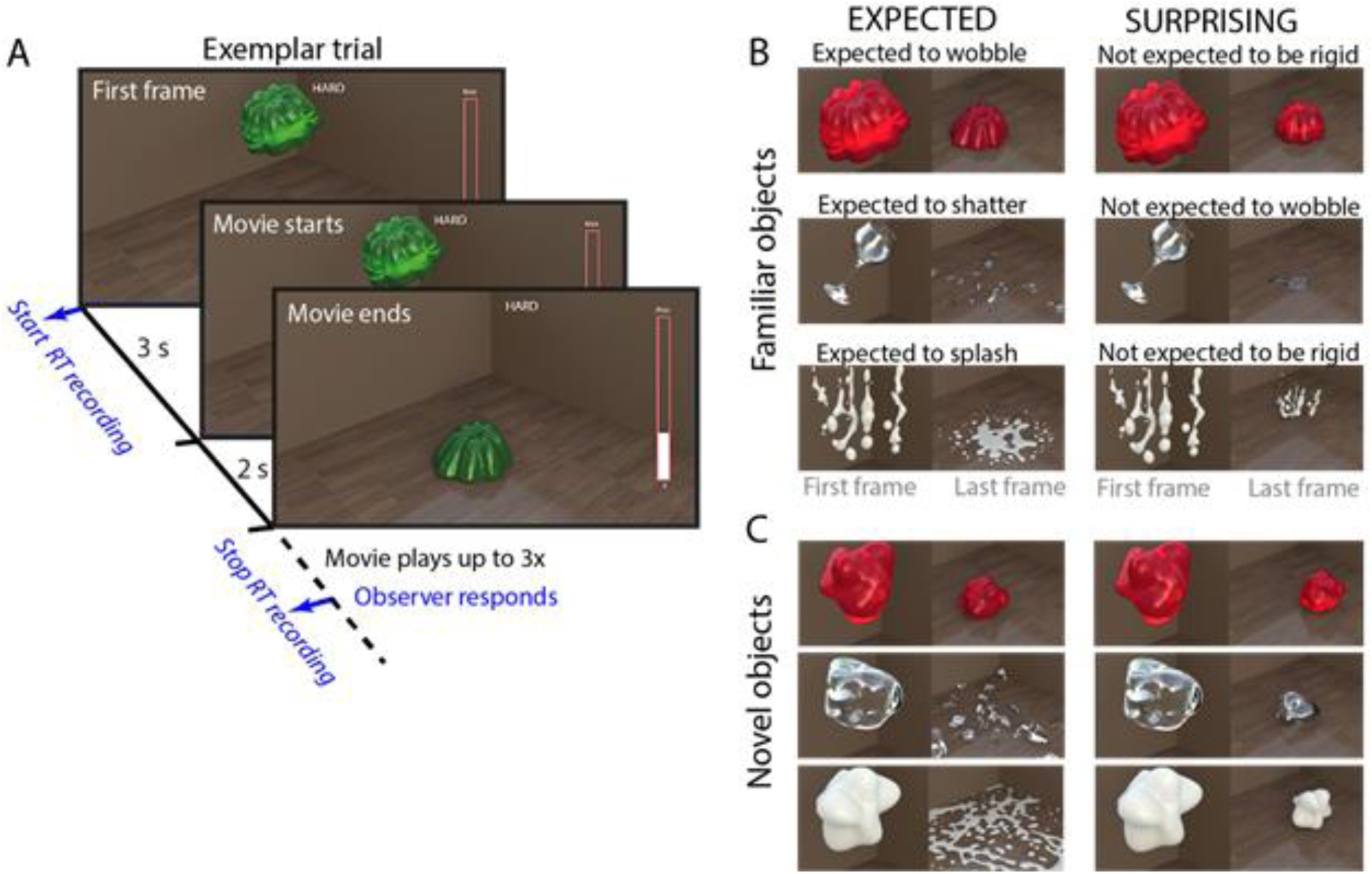
Trial and stimuli. **A.** An exemplar trial. **B.-C.** A subset of familiar objects (B) and corresponding novel objects (C) used in the experiments. Familiar objects could either behave as expected, or in a surprising manner. Note that this distinction (expected vs surprising) is only meaningful for familiar objects. Note that individual scenes are scaled to maximize the view of the object (First frame), or to give a good impression of the material kinematics (Last frame). Figure 3 (Expected condition) and Supplementary Figure 1 (Surprising condition) show corresponding views for the entire stimulus set. The objects were rendered at approximately the same size as each other, so that for example the key was the same “physical” (simulated) size as the chair, even though in real life chairs are larger than keys. We chose to do this so that the objects would hit the ground at the same time, and behave in a similar way under gravity. This is important for some of our analyses (see Analysis - Response time).

#### Animations

Each movie consisted of 48 frames, depicting an object suspended in air, which then fell to the ground. Impact occurred exactly at the 11^th^ frame for all objects. The largest extent of the objects in the first frame varied between 6.91 (clay pot) and 12.6 (spoon) degrees visual angle. The largest extent of the objects in the last frame depended on the deformation, but varied between 48.85 degrees visual angle for shattering/splashing items and 4.29 degrees visual angle for rigidly falling items. Object deformations were simulated using the Rigid Body, Cloth, and Particles System physics engines in Blender. For technical specifications about the Particle System simulations, we refer the reader to the parameters listed in (Schmid & Doerschner, 2018a).

### Apparatus

The experiment was coded in MATLAB 2015a (MathWorks, Natick, MA) using the Psychophysics Toolbox extension (version 3.8.5, Brainard, 1997; Pelli, 1997; Kleiner, M., Brainard, D., Pelli, D., Ingling, A., Murray, R., & Broussard, 2007), and presented on a 24 5/8’’ PVM-2541 Sony (Sony Corporation, Minato, Tokyo, Japan) 10-bit OLED monitor, with a resolution of 1024 x 768 and a refresh rate of 60 Hz. Videos were played at a rate of 24 frames per second. The participants were seated approximately 60 cm from the screen.

### Task and Procedure

Main (Motion) Experiment: Observers were asked to watch a short video clip to the end and then to rate the object they saw on one of four attributes (hardness, gelatinousness, heaviness, and liquidity) as quickly (but as accurately) as they could. We choose the attributes such that they would capture some aspect of the mechanical material qualities of the objects. For example, a splashing object is likely to be rated as very *liquid*, and a non-deforming object not, a wiggling object is likely to be rated as very *wobbly* but a shattering object not. In order to familiarize observers with the rating task, the use of the slider bar and the keypresses, they completed four practice trials with two objects that did not occur in the actual experiment and with two rating adjectives that also did not occur in the experiment (e.g. rate how shiny this object is).

The experiment was organized into four blocks, one block per attribute. Before the block started, the observer was familiarized with the rating question and then proceeded with a button press to start the trials. On every trial, a reminder of the question of this block remained at the top of the screen, e.g. ‘*hard*’ (for ‘How hard is the object?’), together with the first frame of a movie, which was held static for three seconds before the movie was played to the end. After this the movie clip repeated 2 more times (without the hold at the beginning), if necessary. Participants were asked to first watch the video until it finished (i.e. the first play through) and then to rate the object as quickly as they could (while still maintaining accuracy). They indicated their rating by using the mouse to adjust the height of a slider bar placed on the right side of the screen (Figure 4A). A zero setting indicated the absence of an attribute, e.g. not *gelatinous* at all, while a maximum setting would correspond to the subjective maximum value of an attribute. The trial was completed when the observer pressed the space key on the keyboard, after which time the next trial would immediately begin. Response time was measured from the beginning to the end of a trial (between spacebar presses). The slider position of the previous trial was carried over the new trial in order to give the experimental interface a more natural feel to it.

Participants completed 240 trials in total (2 surprise conditions (Expected, Surprising) × 2 object familiarity (Familiar, Novel) × 4 attributes × 15 objects). Surprising condition and object type were the two relevant manipulations in the experiment. While the order of blocks was the same for all observers (*hard, gelatinous, heavy, liquid*), the trial order in each block was randomized.

#### First Frame/Last Frame Experiment

The tasks, setup, and procedures were identical to that of the motion experiment. In contrast to the Motion experiment, in these experiments, the first/last frame of each of the videos was held on-screen for the same duration as a single presentation of the video.

### Participants

#### Motion Experiment

25 participants (mean age 24.8; 18 female) participated in the experiment; 23 were right-handed and all had self-reported normal or corrected-to-normal vision.

#### First Frame Experiment

15 participants (mean age 26.40; 13 female) participated in this experiment; 13 participants were right-handed and all had self-reported normal or corrected-to-normal vision.

#### Last Frame Experiment

14 participants (mean age 26.85; 10 female) participated in the ‘Static Last Frame’ condition of Experiment 1; 12 were right-handed. All participants had self-reported normal or corrected-to-normal vision.

All participants were native German speakers, and the experiment was given entirely in German. The experiment followed the guidelines set forth by the Declaration of Helsinki, and participants were naïve as to the purpose of the experiment. All participants provided written informed consent and were reimbursed at a rate of €8 per hour.

### Analysis

#### Rating differences: Expected versus Surprising

To measure the influence of object knowledge on perceived material properties, we computed rating differences between Expected (Figure 3) and Surprising conditions (Supplementary Figure 1). For each object type (Familiar, Novel), each attribute (*how hard, how gelatinous, how heavy, how liquid*), and each material class (Splashing, Shattering, Non-deforming, Wobbling, Wrinkling), rating differences were calculated between the three objects with Expected outcomes and the three (different) objects with Surprising outcomes (ratings were averaged over the three objects before difference scores were computed). Thus, *how* an object deformed was the same, e.g. it would splash, but *which* object would do the (e.g.) splashing is different in Expected (honey, milk, water) and surprising conditions (chair, pot, curtain).

Our stimulus set was balanced in the sense that for every type of deformation (splashing wrinkling etc.) we found three familiar objects that ‘naturally’ have these material kinematics. In the Surprising condition the deformations were paired with other familiar objects such that a given deformation might be perceived as unusual for this type of object (e.g. a chair splashing). For some of these object-motion pairs it was not clear to predict the particular direction in which the rating will change (e.g. will a Splashing chair look more *liquid*, or *harder*, or *heavier* than Splashing milk?). Therefore, in order to obtain an initial measure of whether expectations affect our perception of object properties at all we took the absolute value of these difference score as an index (Effect of Expectation Index, ***ϵ***), i.e., we were agnostic about the direction of this effect:

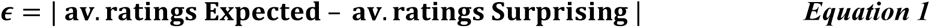

We averaged the ratings in the expected and surprising conditions before finding the difference since it is not clear how to match up objects in the expected and surprise conditions (i.e. whether to match the expectedly rigid chair with the surprisingly rigid curtain or the surprisingly rigid jelly). This resulted in 40 difference scores for each observer, 20 in the Familiar object condition and 20 in the Novel object condition. If object knowledge influences perceived material properties, then ratings should differ between Familiar objects that behave as expected, and Familiar objects that behave in a surprising way, i.e. ***ϵ*** for Familiar objects should be larger than ***ϵ*** for Novel objects (***ϵ*** _Familiar object_ > ***ϵ*** _Novel object_). A binomial sign test was used to compute the likelihood of obtaining *k* or more instances in which ***ϵ*** (averaged across observers) was greater for Familiar versus Novel objects.

#### Prior Pull

The four attributes that observers rated in our experiments are related to the kinematic properties of objects. Such properties can be best estimated when we observe how an object interacts with another one, e.g. if two objects collide and one deforms more than the other, we can tell that one object is softer. Conversely, it should be difficult to judge such properties (hardness, gelatinousness etc.) in static frames, unless we rely on our previous experiences observing how objects interact. Therefore, one could use the First Frame ratings of Familiar objects as a measure of *prior knowledge* about the material qualities from familiar shape and optics associations. Ratings of moving Novel objects, on the other hand, could be used as a measure of how much the image motion, generated by the kinematics of the material (‘sensory’ route), influences the rating (no influence of familiar shape, equating for the effect of familiar optical properties). These two conditions make similar predictions (i.e. yield similar ratings) when the material behavior is expected but make *different* predictions (i.e. yield different ratings) when the material behavior is surprising. We operationally define and measure the “prior pull” as the distance between Familiar and Novel object motion ratings in the direction of First Frame ratings. This could be stated as a conditional statement: a prior pull occurs if:

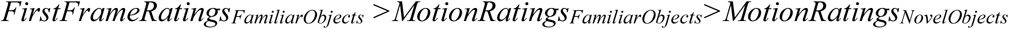

or if

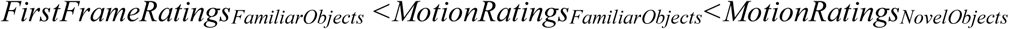

If one of these two conditions is true, then the magnitude of the prior pull can be computed as:

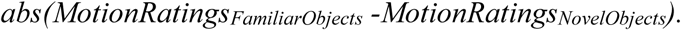

We assess whether prior pull occurs more frequently in the Surprising motion condition compared to the Expected motion condition.

#### Response times

The time taken to make each judgment (response time) was measured. We reasoned that rating the material properties of materials that behave surprisingly might involve the reiterative correction of a prediction error by the visual system, and this error correction might be associated with an increase in response time when rating objects that behave in a surprising way. Before computing the difference in response time between Expected and Surprising conditions, we pre-processed response time data as follows: we subtracted the time to impact (3 seconds static first frame + 0.45 seconds to impact) from the raw response times so that a response time of zero would now indicate time of impact. For each participant, we averaged their response times over objects and attributes. A repeated measures ANOVA was performed on this averaged response time data to determine whether response times differed between expected and surprise conditions, or familiar and novel object conditions.

#### Exclusions

For the first 10 participants in the motion experiment, data points for the Novel object matched to the Surprising key object were excluded due to a stimulus presentation error that showed the rigid version of the object instead of the wobbling one, (10 subjects × 4 attributes = 40 data points excluded). For the remaining 15 subjects this error was fixed.

Data points that were faster than 0.75 seconds after impact (fastest possible button press) were excluded, because a shorter response time than this would indicate that observers started pressing the space bar (for next trial) before watching the impact frame of the movie (which would be a violation of the instructions). Response latencies that were longer than 2 standard deviations above the mean were also excluded. By opting for a 2*SD cutoff as opposed to the traditional cutoff of 3*SD (Magnussen, Idås, & Myhre, 1998) we excluded more of the longer RTs, which occurred more for familiar objects that behaved surprisingly (see Supplementary Figure 2). Therefore, this cutoff criterion was more conservative.

Following these exclusions, approximately 6% of the data were excluded for response times that were too fast or too slow according to this criterium (∼1.3% too fast, ∼4.7% too slow, see Supplementary Figure 2 for a breakdown of exclusions per experimental condition).

## Results

### Ratings and the Effect of Expectation (*ϵ*)

Figure 5 provides an overview of average observer ratings in in all experiments and conditions. Each of the 5 polar plots in each row shows average ratings for 3 (symbolized by icons) of the 15 Familiar objects (circles) and corresponding Novel objects (stars) in response to each of the four questions (*how hard* (red), *gelatinous* (green), *heavy* (magenta), or *liquid* (blue)). The grouping of the objects into one polar plot was done according to how these objects (familiar and corresponding novel) would deform (i.e. non-deforming, wobbling, shattering, wrinkling or splashing). Note, that in the Surprising conditions the object identity and how the object deforms after falling onto the floor mismatch (e.g. shattering water, splashing chairs etc.) Overall, observers’ responses made sense. For example, in Figure 5A static rigid objects were all rated as hard (red cicles), jell-O was rated as very gelatinous (green circle) and liquids were rated as very liquid (blue circles). Ratings for Novel objects (stars) who inherited material properties from corresponding Familiar objects tended to overlap with that of the Familiar objects, however, this overlap tended to vary between the different experiments (Panel A-E): e.g. overlaps in the expected motion condition (Figure 5C) were much larger than in the First frame (Figure 5A) or Last frame conditions (Figure 5B & D), likely owing to the different types of information available in each case (e.g. image motion or not, surprising motion or expected motion etc.). In the regression analysis below, we aim to explain the differences in ratings between Familiar and Novel objects, allowing for different high and low-level factors to account for the data.

**Figure 5.**
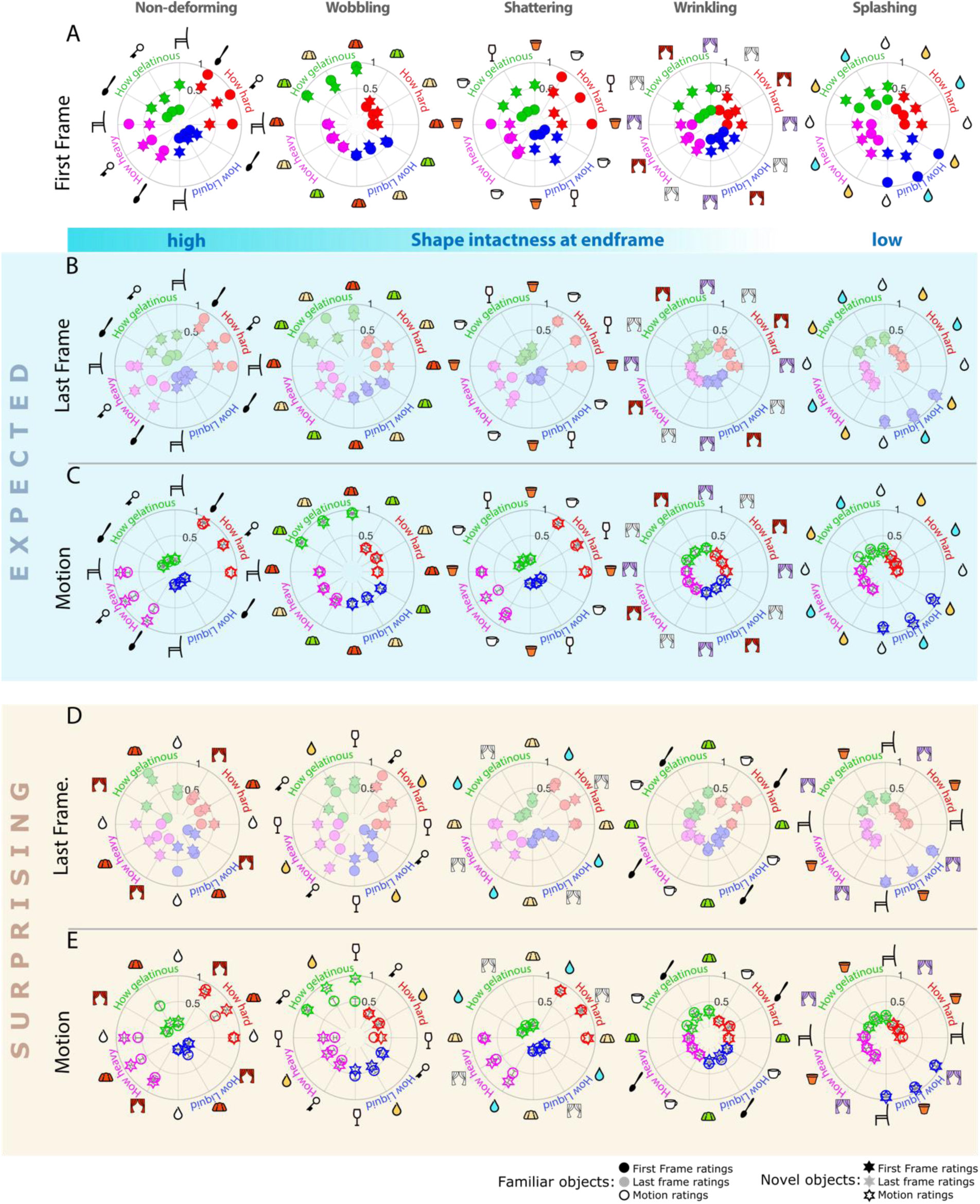
Ratings results from all experiments. Shown are average observer ratings of all experiments for the four different questions about material qualities e.g. how hard, liquid, heavy or gelatinous an object appears. Each column shows the data across experiments for one particular type of material kinematics (wobble, splash etc). Icons symbolize individual Familiar objects (chair, key, cup, pot, glass, spoon, blue droplet: water, yellow droplet: honey, white droplet: milk, violet curtain: silk, red curtain: velvet, white curtain: linen; yellow custard, red and green jell-O). Each question and corresponding data are coded in the same color (red: hard, blue: liquid, purple: heavy, and green: gelatinous). Ratings could vary between 0 (lowest) and 1 (highest). The circle and star symbols correspond to ratings of Familiar and Novel objects, respectively. The symbol style filled, desaturated and open, correspond to the different experiments First Frame, Last Frame and Motion, respectively. Standard deviations denote 1 SE of the mean. Overall, ratings of Familiar and Novel objects tended to overlap much more in the Motion condition (C & E), in particular the expected motion condition (C), than compared to the First Frame (A) and Last Frame (B,D) experiments. In the Surprising conditions the object identity and how the object deforms after falling onto the floor mismatch (e.g. shattering water, splashing chairs etc.). Data from the First Frame (A) and Motion Conditions (C & E) are replotted in Figure 7 in order to illustrate the influence of prior expectation on material quality judgements.

To determine whether object knowledge influences perceived material properties, we computed the Effect of Expectation Index (***ϵ***) for Familiar and Novel objects. We postulated that this effect would be greater for Familiar objects because expectations about material properties should not be triggered as strongly when the object shape is unfamiliar. This is what we found. Figure 6 plots the Effect of Expectation Index (***ϵ***) for each attribute and each material class, averaged across subjects. In all but one condition (19/20), ***ϵ*** was greater for Familiar objects than Novel objects, sign test: p<0.001. Results from a paired *t*-test corroborate the finding that on average, ***ϵ*** was greater for Familiar objects than Novel objects, *t*(24)=6.70, *p*<.001, which also verifies the effect at an individual level. This supports our hypothesis that judgments of material qualities are not based purely on the observed material mechanics, but are also affected by prior knowledge about the typical material mechanics of a (familiar) object. Note that the effect of expectation index for Novel objects was not zero, which suggests that observers might generate some predictions about the mechanical properties based on the estimated optical properties of these objects, and/or based on associations they have with the general material properties of a bounded convex shape, e.g. a rigidity prior (Ullman, 1976; Grzywacz & Hildreth, 1987; but also see Jain & Zaidi, 2011). Below we break down these specific influences on material judgements and based on this develop a linear regression model that tries to account for the observed rating differences in Expected and Surprising conditions.

**Figure 6.**
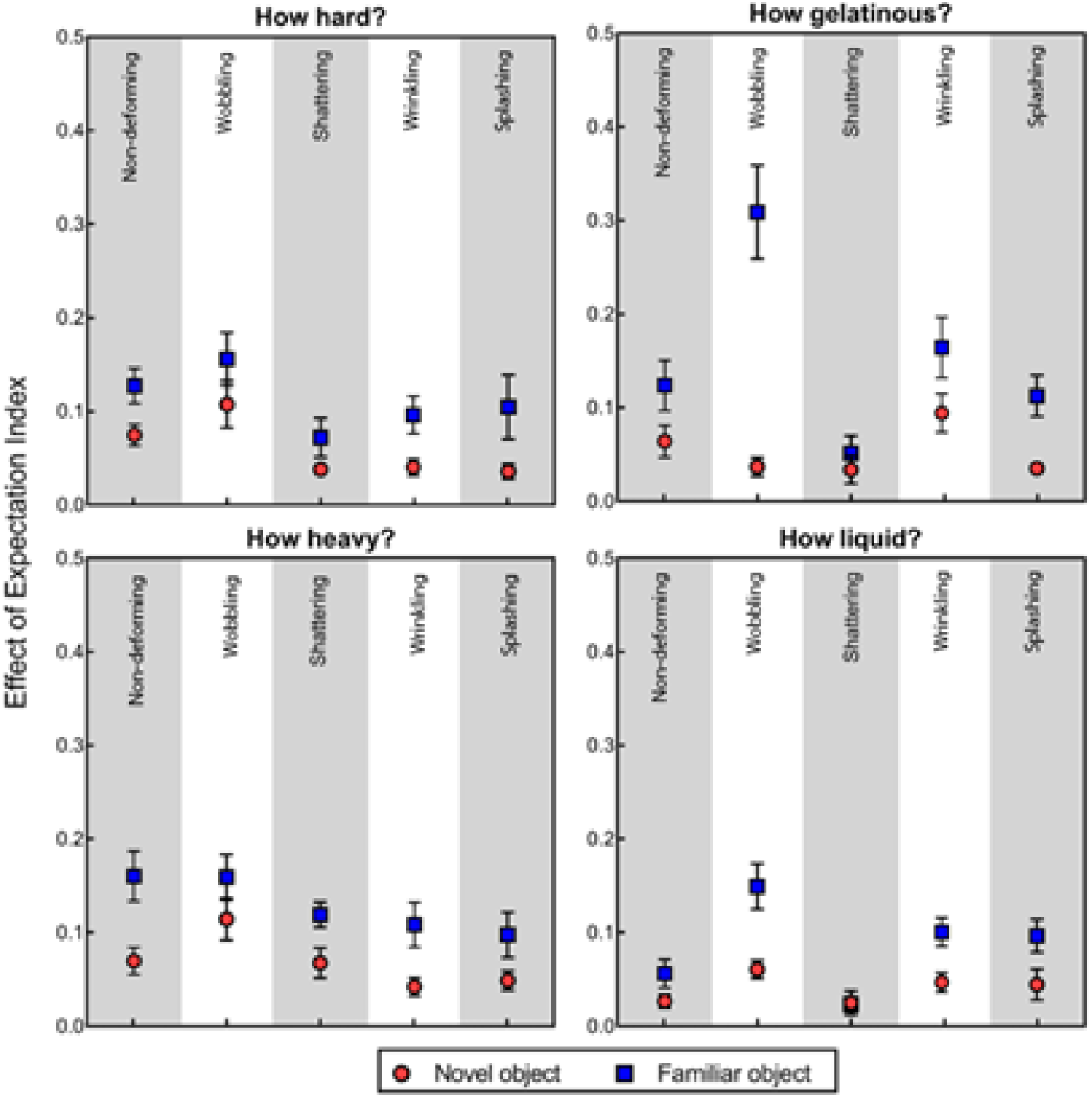
Effect of Expectation Index (*ϵ*) averaged across subjects. **ϵ** was calculated as the absolute difference between ratings in the Expected and Surprising conditions (see analysis section, Equation 1). Error bars are one standard error of the mean.

### Prior Pull

In some cases, the directionality of the rating differences (Expected vs Surprising) is directly interpretable. For example, the spoon that wrinkled surprisingly (Figure 5E, fourth plot) was rated as *harder* (red circle, .25) than the linen, silk, or velvet curtains that wrinkled expectedly (Figure 5C, .18). Prior knowledge about spoons being hard seems to have led to increased ratings of *hardness* compared to their soft curtain counterparts, despite all of these objects wrinkling. Thus, our results suggest that prior object knowledge about *hardness* “pulled” ratings of *hardness* towards this expectation. We visualize this ‘prior pull’ in the data in Figure 7.

**Figure 7.**
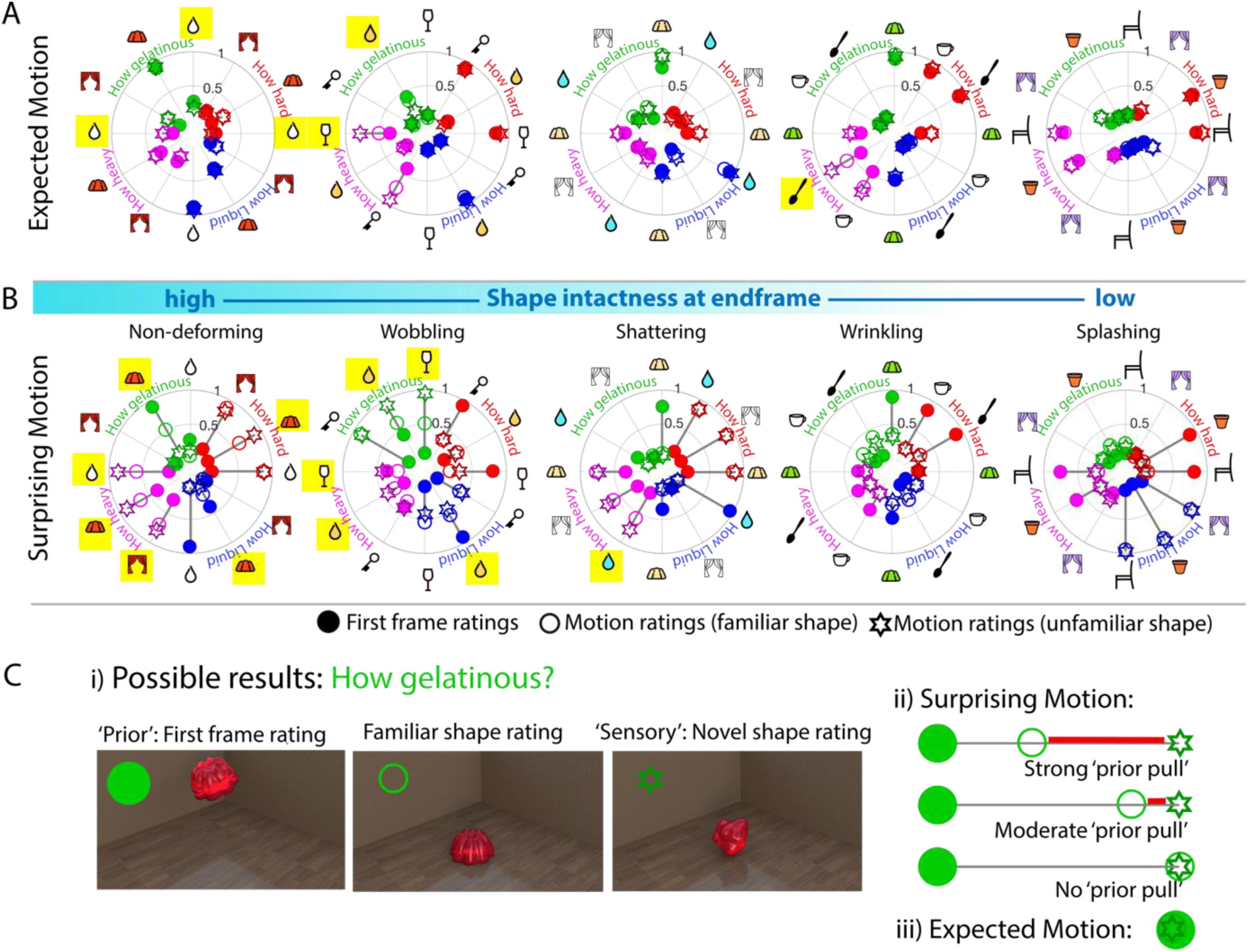
Material quality ratings and prior attraction. Results of the First frame and Motion conditions from Figure 5 are re-plotted, keeping the same symbols and notation. A. Average observer ratings from 3 conditions (i.e. First Frame Familiar objects (filled circles), typically-behaving Familiar objects (unfilled circles) and corresponding (moving) Novel objects (unfilled stars)) tended to overlap. The difference between First frame ratings for Familiar objects and ratings of moving Novel objects is indicated by a dark grey line. The organization of objects follows that in panel B. B. Same as A, but here, ratings of atypically-behaving familiar objects are plotted as unfilled circles (organized by type of motion), and ratings of corresponding Novel objects - i.e. unfamiliar shapes-- which inherit their optical and kinematic qualities from a familiar object - as unfilled stars. The motions are arranged according to how much the object remains intact and recognizable after impact on the floor (also see Supplementary Figure 3). Yellow highlighted symbols show a statistically significant prior pull. See main text for more detail. The yellow highlighted cases show that “prior pull” occurred more in conditions where the object was still intact and recognizable at the end of the movie (objects that behaved rigidly or wobbled). Supplementary Table 1 lists corresponding statistics and p-values. C. i) Illustrates how we measure how much the rating of an atypically-moving Familiar object (middle image) overlaps with the rating of a material-matched moving Novel object (right image), or conversely, how much it is pulled towards ratings of a static view of the Familiar object (left image). ii) Shows possible results: For example, seeing an image of red jell-O in its classical shape, observers tend to expect that it is quite gelatinous. When they see an object with the same optical properties that falls and does not wobble when it hits the floor, they rate it as very non-gelatinous, i.e. we have a large rating difference (gray line). When a classically-shaped red jell-O falls on the floor and doesn’t wobble, observers could either rate it similar to the novel object -after all it doesn’t wobble at all (no prior pull) - or it could be rated as somewhat more gelatinous, despite the sensory input, possibly because prior experience influences the appearance, making observers perceive wobble when there isn’t (prior pull, red line). iii) When the familiar object moves exactly as expected, and when there is no strong influence of shape familiarity on material judgements, all three ratings will overlap.

Prior pull can only occur if ratings on a given quality in First frame (filled dots) and Surprising motion (open stars) conditions differ substantially, as indicated by the length of the dark gray lines between these two types of data points in Figure 7. We would expect to find this occurring more frequently in the Surprising motion condition, and our results confirm this: First Frame ratings differed significantly from Novel object motion ratings for 48 out of 60 (80%) conditions in the Surprising motion condition, and only 24 out of 60 (40%) conditions in the Expected motion condition (also see Table 1). A meaningful prior ‘pull’ occurs only when the ratings of atypically-moving (Surprising) Familiar (unfilled circles) and Novel objects (unfilled stars) do not overlap and when ratings of atypically-moving (Surprising) Familiar objects are pulled in the direction of First frame ratings (filled circles; see Figure 7C, ii). Significant cases of the prior pull effect are highlighted in yellow in Figure 7 (see Supplementary Table 1 for significance tests). If, instead, ratings of moving Familiar (open circles) and Novel objects (open stars) overlap completely, this shows that the object shape prior did not exert any significant influence on the rating (see Figure 7C, ii). We found that the more the familiar object remained intact and thus recognizable in the Surprising motion condition (Figure 7B, left plots), the more likely the prior exerted an influence over the material appearance (also see Supplementary Figure 3, which assesses object intactness).

That only a subset of our objects exhibits a “pull” towards the prior is consistent with literature that shows when sensory input is unambiguous, it will dominate the percept, and prior knowledge does not have an effect (Summerfield & De Lange, 2014). In our case, particular combinations of shape and optics (often involving “shape recognizability”) at the end of the animation may lead observers to essentially “ignore” sensory information. For example, the rigid jell-O is perceived as gelatinous despite not wobbling. Interestingly, prior associations with specific combinations of motion, shape, and optics at the end of the movie may “enhance” differences between Familiar and Novel object ratings (e.g. in the case of the wobbling glass, the particular combination of translucent optics forming what looks like a puddle might evoke strong representations of liquidity, so despite the wobble, it is not considered gelatinous).

One might argue that the reason why we see relatively few instances of significant prior pull is that Novel objects might also elicit expectations (after all, they are bounded shapes with particular optical properties). The correlations in Figure 8 between First Frame and motion ratings, however, suggests that, for the most part, this is not the case: while Familiar object first frame ratings predicted Expected motion ratings extremely well (Figure 8A, R^2^=0.9), Novel object first frame ratings predict motion ratings very poorly (Figure 8B, R^2^=0.07). These results are also supported by the average interobserver correlations in Figure 8C. Interobserver correlation is much lower for novel objects in the First frame experiment than for familiar objects suggesting that static images of Novel objects do not elicit consistent (typical) expectations about kinematic material properties across observers. Thus, for the most part, Familiar object first frame ratings are a good measure of observers’ prior expectations about the material qualities of these objects. In contrast, Novel object motion ratings are an ideal measure of sensory input because the motion from the material behavior includes information about both the mechanical deformations, as well as any additional effect of image motion from optics (Doerschner et al., 2011).

**Figure 8.**
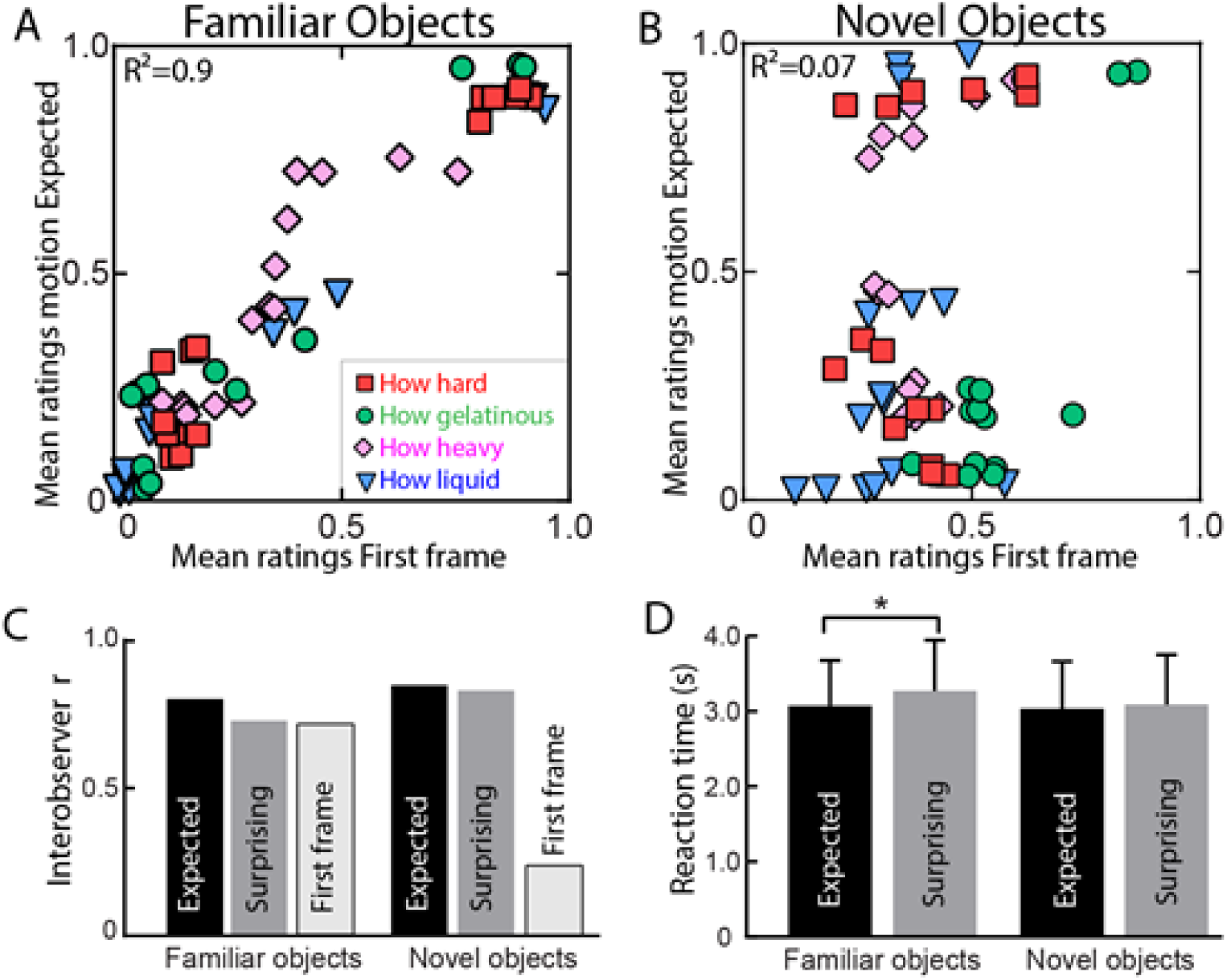
Prediction strength response time differences and interobserver correlations. **A.** Correlation between mean First frame ratings and mean Expected motion ratings for Familiar and Novel objects (**B.**) A high correlation indicates that the First frames (still images) of objects are highly predictive of the objects’ kinematic properties, and thus are in good agreement with ratings in the Expected motion condition, where objects fall and deform according to their typical material kinematics. This is clearly not the case for Novel objects, suggesting that these objects do not elicit strong prior expectations about how an object will deform. **C.** Average interobserver correlation for Expected and Surprising motion trials, as well as the First frame experiment. Note that only for Novel objects, this latter correlation was quite low, suggesting that still images of unfamiliar objects do not elicit a strong prior in observers about the material qualities measured in this experiment. **D.** Response time data averaged across all observers for Expected (black) and Surprising trials (medium gray). Stars indicate significant differences p<0.001. Error bars are 1 standard error and show variability between subjects.

### Linear Regression Models

Given that there are a few cases where Novel objects do seem to generate correct predictions about the material outcome (those at the bottom left and top right of Figure 8B), and since some of the magnitude of the prior pull may be explained by shape recognizability at the end of the movie, we tested a linear regression model that predicted the direction and magnitude of Familiar and Novel object rating differences from prior pull from optics, familiar shape, and shape recognizability at the end of the animation (Figure 9). We assumed that Novel objects would have less pronounced prior associations and this alone can well account for the rating differences we observed. From our data we modelled 3 different association routes that could potentially influence the rating: optics-associations (H1), shape-associations at the beginning of the movie (H2), and shape-associations at the end of the movie (H3).

**Figure 9.**
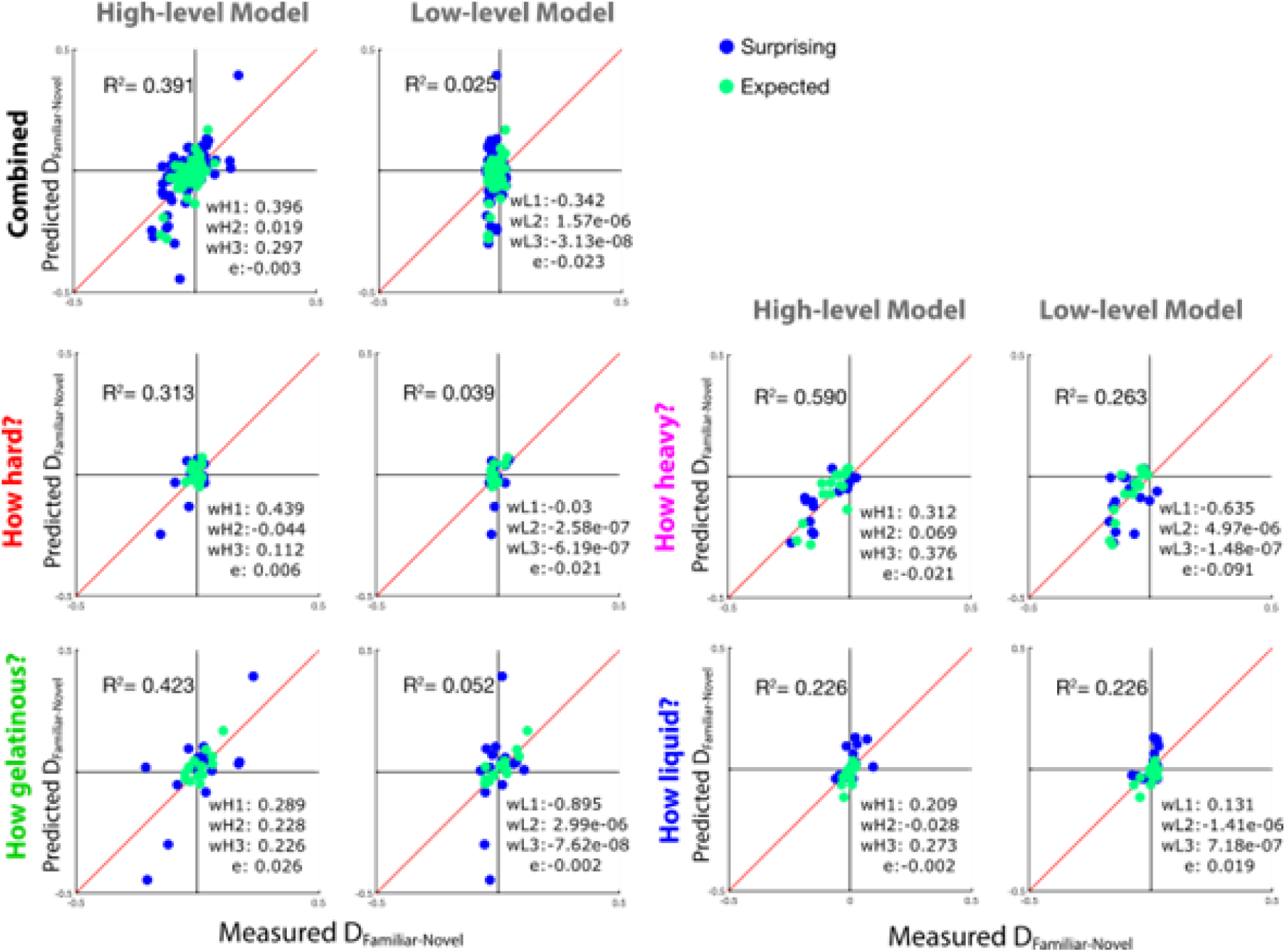
Low and high level linear regression models predicting the difference between ratings of moving familiar and control objects D_familiar-novel_. We developed two models with the aim to account for the differences we observed in ratings of familiar and novel moving objects. The computation of the individual predictors is described in the main text (analyses section). In the lower right inset of each plot we show the weights (w) of each predictor (H1: founded shape & optics prior, H2: shape familiarity, H3: last frame shape recognizability, L1: motion energy difference (also see Supplementary Figure 5), L2: object size differences first frame, L3: object size differences last frame). Overall, the high-level model was more successful in predicting this difference than the low-level model (top two panels: ‘combined’). However, this pattern varied as a function of rating questions: the high-level model performed best for ratings of gelatinousness and hardness, whereas the low-level model performed as good as the high-level model for ratings of liquidness. The latter is likely due to the fact that ratings of liquidness might be strongly modulated by how much a substance physically spreads in the image.

Each predictor in the high-level model was multiplied by a weight G that took into account the predictability of the “behavior” of a given stimulus (computed as the difference between first frame and motion ratings in the Expected condition) and the extremeness of the rating (computed as the twice the difference between first frame ratings in the Expected condition and .5). Thus, predictors in the high-level model were:

H1 (=H1*G), where G= predictability*extremeness: the optics & bounded shape prior (computed as the difference between First Frame ratings of Novel objects in the Expected condition and moving stimuli ratings of Novel objects in the Expected condition). Rationale: This predictor captures to what extent the reflectance properties and the fact that observers see a bounded shape will lead to prior expectations about the material kinematics.

H2 (=H2*(1-G)): shape familiarity (computed as the overall differences between First frame ratings of Familiar objects and corresponding ratings of moving Novel objects (across both expected and surprising conditions). Rationale: This predictor captures to what extent expectations generated by the familiar shape of the object can explain the rating differences, not controlling for the effects of optics and bounded shape (which are captured by H1).

H3 (=H3*(1-G)): last frame shape recognizability (computed as the overall differences between Last frame ratings of Familiar and Novel objects (across both, expected and surprising conditions). Rationale: The reason for including this predictor is our hypothesis that the conflict between prior and surprising motion should be larger, if the object would be still recognizable after falling to the ground (as in the wobbling or rigid motion conditions, see Table 1.)’

We can model the data moderately well with this *high-level* model (see analysis, R^2^ between 0.266 and 0.59 depending on the question), but not perfectly, potentially due to specific motion-shape-optics interactions (Schmid & Doerschner, 2018).

Although retinal size was approximately equated between novel and familiar objects, their area and 3D volume was not. Moreover, shape and volumes within familiar object stimuli varied quite a bit. A voluminous sphere-like object will behave quite differently than a spoon (imagine how much each of them would splash or wobble upon impact if made from liquid or jelly). Therefore, such stimulus variations might to some degree account for rating differences between the familiar and corresponding novel objects and might thus provide alternative explanations to our ‘effect of expectation’ hypothesis. In order to test the degree to which such low-level effects account for the rating data we developed a low-level regression model, which contained 3 predictors that aimed to capture the most salient low-level differences between stimuli.

L1: motion energy difference (this was computed in two steps): 1. Take the sum of the absolute value of all consecutive image differences, starting with the impact frame, e.g. sum(abs(f11-f12), abs(f12-f13), …, abs(f39-f40)), (Tschacher, Rees, & Ramseyer, 2014), for all experimental conditions/stimuli (60 in total) and normalize these values by object size (number of pixels corresponding to the object on frame 11 MEn), 2. For each object in the expected and surprising conditions, compute the differences of MEFamiliar and MENovel conditions. Rationale: Familiar and Novel objects were often different in 3D volume and shape, thus differences in ratings might be attributable to resulting differences in motion energy when these two classes of objects deform.

L2: first frame object size (computed as number of pixels in the first frame corresponding to the object). Rationale: Differences in the area taken up by Familiar and Novel objects in the first frame might account for rating differences, e.g. if the Familiar object was smaller, it might have generated predictions to be less heavy than the corresponding Novel object (irrespective of the familiar shape of the object).

L3: last frame object size (computed as numbers of pixels in the last frame corresponding to the object). Rationale: this predictor is related to H3 in the high-level model (below). Differences in the area taken up by Familiar and Novel objects in the last frame might account for rating differences, e.g. if the novel object ‘splashed’ more (thus took up more area in the last frame) it might have been rated more liquid simply due to this fact.

Such a model performs extremely poorly (R^2^ = 0.025, p>0.05, Figure 9). This suggests rating differences are not due to differences in object size or image motion.

### Response times

We find a small but significant increase in RT in the Familiar object surprising condition - which is the condition that most strongly juxtaposes prior expectation with sensory evidence (Figure 8D). There was no significant response time difference between expected and surprising Novel objects conditions. This was assessed with a a two-way repeated measures ANOVA, which revealed a main effect of object familiarity (participants took longer to respond to familiar versus objects), F(1,24)=14.72, p=0.0008, and a main effect of surprise (participants took longer to respond to surprising versus expected events), F(1,24)=13.24, p=0.013. However, these main effects must be interpreted in light of the significant interaction between object familiarity and surprisingness, F(1,24) = 6.395, p=0.0184. Follow-up t-tests (Bonferroni corrected) show that participants took longer to respond to surprising versus expected stimuli when objects were familiar, t(24) = 4.911, p<0.0001. However, this difference was not significant for novel objects, t(24) = 1.334, p=0.1946.

## DISCUSSION

Visual perception is not a one-way (bottom-up) road; how we process visual input is influenced by expectations about the sensory environment, which develop from our previous experience and learning about existing regularities in the world, i.e. associating things or events that co-occur. Expectations have been shown to facilitate visual processing in the case of priming, to modulate the frequency of a particular percept in bi-stable stimuli, and to change our interpretation of ambiguous stimuli (see Kveraga et al., 2007, or Panichello, Cheung, & Bar, 2013 for a review). However, the stimuli used in these experiments have been fairly simple (static images of objects), and it has been shown that learning associations can also include fairly complex phenomena. For example, recently, Bates, Yildirim, Tenenbaum, & Battaglia, (2015) (also Kubricht et al., 2016; Battaglia, Hamrick, & Tenenbaum, 2013), showed that humans can learn to predict how different liquids flow around solid obstacles (also see other examples for predicting of motion trajectories of rigid objects, e.g. (Flombaum, Scholl, & Santos, 2009; Gao & Scholl, 2010, or Soechting, Juveli, & Rao, 2009). While the authors attributed human performance to an ability to “reason” about fluid dynamics, here we explicitly test whether existing *perceptual expectations* about material properties can set up rather complex predictions about future states, and whether – and to what extent – these expectations influence material appearance. We show that the qualities of “surprising” materials (Itti & Baldi, 2009a) are perceived different to expected ones that behave the same (Figure 7), and that surprise leads to increases in processing time of the stimuli (Figure 8D, in line with Itti & Baldi, 2009b; Baldi & Itti, 2010; or recently Urgen & Boyaci, 2019). Our method provides a general technique to differentiate the extent to which material qualities are directly estimated from material kinematics versus being modulated by prior associations from familiar shape and optical properties, and contributes thus an essential piece to the puzzle of how the human visual system accomplishes material perception (Adelson, 2001; Maloney & Brainard, 2010; Anderson, 2011; Fleming, Nishida, & Gegenfurtner, 2015; Fleming, 2014; Komatsu & Goda, 2018)

### Knowledge affects (material) perception

There are countless demonstrations showing that knowledge affects how we perceive the world: from detecting the Dalmatian amidst black and white blotches, or identifying an animal in the scene (Thorpe, Fize, & Marlot, 1996), deciding on the identity of a blob (Oliva & Torralba, 2007), or being a greeble or bird expert (Gauthier, James, Curby, & Tarr, 2003), knowledge directly influences our ability to perform in these instances. Object knowledge does not just facilitate categorical judgments, it also affects the estimation of visual properties such as color or motion (Hansen et al., 2006; Olkkonen & Allred, 2014; Scocchia et al., 2013). How exactly knowledge alters and facilitates neural processes in visual perception is a topic of ongoing research (e.g. Rahman & Sommer, 2008; Gauthier, Skudlarski, Gore, & Anderson, 2000; or Kveraga et al., 2007).

In contrast, the role of predictions or associative mechanisms in material perception is not well understood (van Assen et al., 2018; Schmidt et al., 2017; Paulun et al., 2017). Knowledge about materials entails several dimensions and can include taxonomic relations: gold is a metal, metals are elements with physical properties, metals are usually malleable and ductile. These classes of metal also have their own perceptual regularities: gold looks yellowish, often has a very shiny, polished and smooth appearance, feels cool to the touch, etc. Our experimental results suggest that identifying a material (i.e. knowing what it is) not only co-activates its typical optical qualities, but also elicits strong predictions about the typical kinematic properties and resulting material ‘behaviors’. For example, liquids are not only translucent or transparent, they also tend to run down, splash, or ooze. Importantly, we seem to have quite specific ideas of what running down, splashing, or oozing should look like (e.g. Dövencioglu, van Doorn, Koenderink, & Doerschner, 2018 probed such ‘ideas’ explicitly), supposedly because we have ample visual (but also haptic) experiences with liquids, and thus opportunities to learn the regularities (statistical or other) associated with a specific material category. Our results show that these specific ideas, or ‘priors’, about material behaviors interfere with the bottom-up processing of visual information, leading to predictable differences in ratings of material properties between Expected and Surprising conditions.

Interestingly, Sharan, Rosenholtz & Adelson (2013) measured reaction times in a material categorization task while participants judged objects made from ‘real’ (e.g. a cupcake made from dough) and ‘fake’ materials (e.g. a knitted cupcake figure). They found a substantial decrease in reaction times at very short presentation times in the ‘real’ condition. Given our results that show response time increases in the surprise condition for Familiar objects, it would be interesting to know if their decrease is primarily driven by an increase the surprising stimuli in their experiment, i.e. ‘fake’ ones – since for those, the object shape/identity (a natural/edible thing) was in conflict with the material of the object (plastic/inedible).

### Prior pull

The yellow highlighted cases in Figure 7 show that “prior pull” occurred twice as much in the Surprising than the Expected condition, and more in conditions where the object was still intact and recognizable at the end of the movie (objects that behaved rigidly or wobbled). Prior pull in the Expected condition also occurred where estimation (sensory input) and associative (prior knowledge) accounts made different predictions (gelatinousness of the honey, heaviness of the wine glass, Figure 7A). In these cases, the ‘expected’ cases were not so expected – this may be related to shape properties, or the size of the splashing of the liquids. Although we controlled for the effects of image motion from optics (e.g. specular highlights), perhaps other low-level image differences exist between Familiar and Novel objects that could be driving differences in ratings. However, we were able to rule this out, since a linear regression model that incorporates these low-level effects as predictors performs extremely poorly. This suggests rating differences were not due to differences in object size or image motion.

We did not aim to test an exhaustive list of material attributes, but to determine whether effects of prior associations on visual input might depend on the type of material attribute judged, and on how the object behaves under external forces. We found that “mechanical” qualities like hardness and liquidity appear to be more directly estimated from material kinematics in “shape destroying” conditions (splashing, shattering, wrinkling), but prior associations play a modulatory role (to the extent where material kinematics can even be ignored (e.g. red and green jell-O) when shape remains somewhat intact. These latter conditions seem to create more of a cue conflict, and are more ambiguous. On the other hand, qualities like gelatinousness and heaviness (which are much more difficult to estimate directly from mechanical deformations) were more affected by familiar shape and optics associations.

### Alternative explanations for rating differences

One might argue that the ‘prior pull’ we demonstrated here is not perceptual but in fact is due to a particular ‘cognitive strategy’ of some observers (i.e. explicitly ignoring the motion information and thus rating material qualities of atypically moving familiar objects as they rated objects on the first frame, while other observers’ ratings were 100% identical to novel object ratings). This would have resulted in bimodal rating distributions and/or low interobserver correlation in the object motion condition, neither of which we found (Supplementary Figure 4 and Figure 8C, respectively).

Another argument against the cognitive strategy approach is supported by response time (RT) patterns in our experiments. A small but significant increase in RT in the familiar object surprising condition - which is the condition that most strongly juxtaposes prior expectation with sensory evidence - would be consistent with the idea of recurrent prediction error correction (Urgen & Boyaci, 2019, Figure 8D). Importantly, we do not find evidence for a response time advantage in expected familiar objects condition (compared to novel, also consistent with (Urgen & Boyaci, 2019), which suggests that this increase in RT cannot simply be due to the fact that observers positioned the slider in advance to the ‘wrong’ position, i.e. to a position that would be more consistent with a rating based on the First frame information only. If observers adopted such a strategy, we should have also seen faster RTs for expectedly moving familiar objects.

Although we do not believe that cognitive strategies were driving our results per se, we do acknowledge that it is unlikely that ratings tasks directly measure the “appearance” of materials, as it is not known what the relevant perceptual dimensions of visual experience are. Finding the relevant perceptual dimensions of materials is an active area of research (e.g. Schmid & Doerschner, 2018, Toscani, Yücel, & Doerschner, 2019). Many studies have investigated the visual perception of properties like softness or gloss (see Fleming, 2017, for a review), and we believe our participants’ ratings in our study results reflect perception to the same degree as these other studies.

It is quite striking that the same material deformations were rated differently in the Expected and Surprising conditions. Toscani, Valsecchi, & Gegenfurtner (2013) showed that depending on the task, observers pointed their gaze at specific points on the stimulus, e.g. near the brightest regions on an object for lightness judgements. One possibility could thus be that the task, object knowledge, and expectations about the material ‘behavior’ guided eye movements of observers in our experiment to specific locations on the stimulus. While in Expected conditions fixation patterns might have been optimal with respect to the task, e.g. observers correctly anticipated how the object would deform (or shatter, splash, wobble etc.), it is possible that in the Surprising conditions the ‘wrong’ expectation guided eye movements to the ‘wrong’ locations on the object, which in turn lead to a different sampling of information and ultimately influenced their judgements of material qualities. We are currently investigating this possibility directly.

### A Bayesian account of rating and reaction time differences

We believe that the results of this study fit well within the Bayesian framework, which offers an account of how prior knowledge is integrated with sensory input. Our experiment constitutes a situation not unlike classical cue-conflict experiments (e.g. Ernst & Banks, 2002; Knill, 2007), where the sensory cues may be in conflict with one another and/or the prior belief. While we do not aim to model our results formally, we still believe that this analogy is useful in interpreting our findings. We will first focus on the rating differences that we found in the Expected and Surprising conditions. In the latter, in many instances, ratings were ‘pulled’ towards the expected material property, not the signaled one. For example, a wrinkling spoon was rated harder than any of the wrinkling control objects. In this latter example, we have 2 cues to object hardness: object shape (a spoon) and the object behavior (it is wrinkling). Here, the cues to hardness are in conflict, the (familiar) shape of the object (in the first frame, which observers saw for 3 seconds) suggest a very hard object, while the subsequent motion information when the spoon impacts suggests a very soft material. In this situation the visual system has two possibilities: 1: in light of the strong sensory (i.e. image motion) information, it could simply completely reject the idea that this object has any degree of hardness and veto the familiar shape cue completely, which classic cue combination would suggest (e.g. outlier rejection (Landy et al., 1995)). As a result, we would see no difference between wrinkling spoon, wrinkling curtains or wrinkling novel objects. 2. the visual system entertains multiple priors (strong and weaker ones) about the state of the world, and that, depending on the sensory input, it adjusts the weights of these priors (Knill, 2007). In our concrete example this would mean that the visual system may entertain multiple priors on spoon hardness - say from softish to very hard, and that instead of vetoing the idea that the spoon was ever hard, the hardness-cue from familiar-shape is only weighed down and integrated with other available cues (e.g. image motion information). This would yield a hardness rating different from that of the wrinkling control objects, or curtains, and this is what we see in our data.

Thus, the robust cue integration model seems to offer a good explanation for the differences in rating data between novel and familiar objects in the Surprise condition. This model was originally developed by Knill (2007) to account for observer behavior in situations with large cue-conflicts, in which our experiment clearly is. However, it seems to only account well for the data when the shape of the object is still recognizable towards the end of the animation (Figure 7, e.g. non-deforming, or wobbling in the Surprise condition). When instead the shape is unrecognizable upon impact (e.g. when splashing), we see more of a situation analogous to cue-vetoing, i.e. there is no difference in perceived material quality between familiar and novel object ratings. However, a ‘destruction’ of the shape has made the (familiar) shape cue completely unavailable which might have changed the integration situation entirely, which makes an interpretation in these cases difficult.

When faced with violated expectations, as in our Surprising condition, the visual system needs to update the generative model in order to minimize the prediction error i.e. the error between the expected state and ‘measured’ state of the world, in order to perform future tasks (Urgen & Boyaci, 2019). Because this updating is a reiterative process, we reasoned that it would take observers longer to perform perceptual tasks when judging material attributes of surprisingly deforming objects. This is exactly what we found. One might criticize that response times for ratings were much longer than times measured in classic reaction times studies (e.g see a review by Eckstein, 2011 on visual search). However, it is not all that uncommon to consider response times of 2 seconds and longer, as in categorical color perception (Boynton & Olson, 1990; or Okazawa, Koida, & Komatsu, 2011). Note that we treated response time data as conservatively as classic reaction time studies, e.g. by removing data points that were two standard deviations above the mean.

Our study bears resemblance to the work by Fujisaki et al. who investigated how different kinds of information sources, namely visual and auditory, are combined in material categorization and material property rating tasks. Some of their stimulus conditions were not unlike our cue conflict scenarios (e.g. combining a visual glass stimulus with a bell pepper sound). They found that for the material rating task, the integration of the two types of information follows a weighted average rule - were the weights depend on the reliability of the respective signals. This reliability was in part related to the task: for example, participants gave higher weights to visual cues for judgements of color and gave higher weights to auditory cues for judgments of pitch or hardness. Also in our results in Figure 9 we see systematic changes in the regression weights of both, high- and low-level models as a function of rating task: for example in the low-level model, the visual cue “motion energy” (wL1) receives more weight explaining rating differences between familiar and novel objects than for ratings of e.g. heaviness, or similarly in the high-level model: the predictor “shape familiarity” (wH2) played a much larger role for explaining rating differences between familiar and novel objects for ratings of gelatinousness that for ratings of liquidness. In order to determine whether this shift in weights is indeed associated with the reliability of the respective cues requires further experimentation, where cue reliability would be manipulated directly.

### Estimation vs Association

Our work shows that previously acquired object-material associations play a central role in material perception and are much more sophisticated than previously appreciated. Previously there have been conflicting findings in the literature about the relative influence of optics, shape, and motion cues to the perception of material properties (e.g. van Assen & Fleming, 2016, Aliaga, O’sullivan, Gutierrez, & Tamstorf, 2018, Schmid & Doerschner, 2018). For example, while some work proposes that perceived material qualities like softness are strongly influenced by motion and shape cues, which completely dominate optical cues (Paulun, Schmidt, van Assen, & Fleming, 2017b), other work showed that both optical and mechanical cues affect estimates of softness (Schmidt, Paulun, van Assen, & Fleming, 2017). Schmidt et al. (2017) suggested that shape recognisability after deformation (e.g. recognising that an object used to be a cube in Paulun et al.’s study) affects how reliable shape cues versus surface optical cues are when judging material properties, thus leading to shape cues dominating over optical cues in Paulun et al.’s study. Our results back up the idea that familiar shape affects observers’ ratings of material properties. The interactions between familiar shape, optical and motion properties is something that future material perception studies should consider and investigate further.

This study not only shows that existing perceptual expectations about material properties can set up rather complex predictions about future states of materials, it also extends a growing theme in the material perception literature that studying the perception of kinematic material qualities can serve as a tool to guide investigations of the neural mechanisms about material properties, as it provides insight into components (high and low level) that make up material perception as a whole (Schmid & Doerschner, 2019).

## Conclusion

This work shows that the visual system can predict the future states of rigid and non-rigid materials. Such predictions can be activated by the shape of an object and – to a lesser extent - also by the optical qualities of a surface. Understanding how high-level expectations are integrated with incoming sensory evidence is an essential step towards understanding how the human visual system accomplishes material perception.

## Contributions

LA, KD, and AS conceptualized and developed the experiments in this paper. AS created the stimuli and programmed the experiments. AS, LA, and KD developed and performed the analyses. KD, AS, and LA wrote the paper.

## Acknowledgements

This work has been supported by a Sofja Kovalevskaja Award endowed by the German Federal Ministry of Education. We thank Roland Fleming, Huseyin Boyaci, and Karl Gegenfurtner for helpful discussions and feedback on earlier versions of this manuscript, and Flip Phillips for providing the ‘Glaven’ 3D meshes.

## Notes

https://www.dropbox.com/sh/bmlf4z03mgfqfna/AABUSwubHleEHp_l1lYkgKkEa?dl=0

### Competing Interest Statement

The authors have declared no competing interest.

