## Supplementary material for "Expectations affect the perception of material properties"

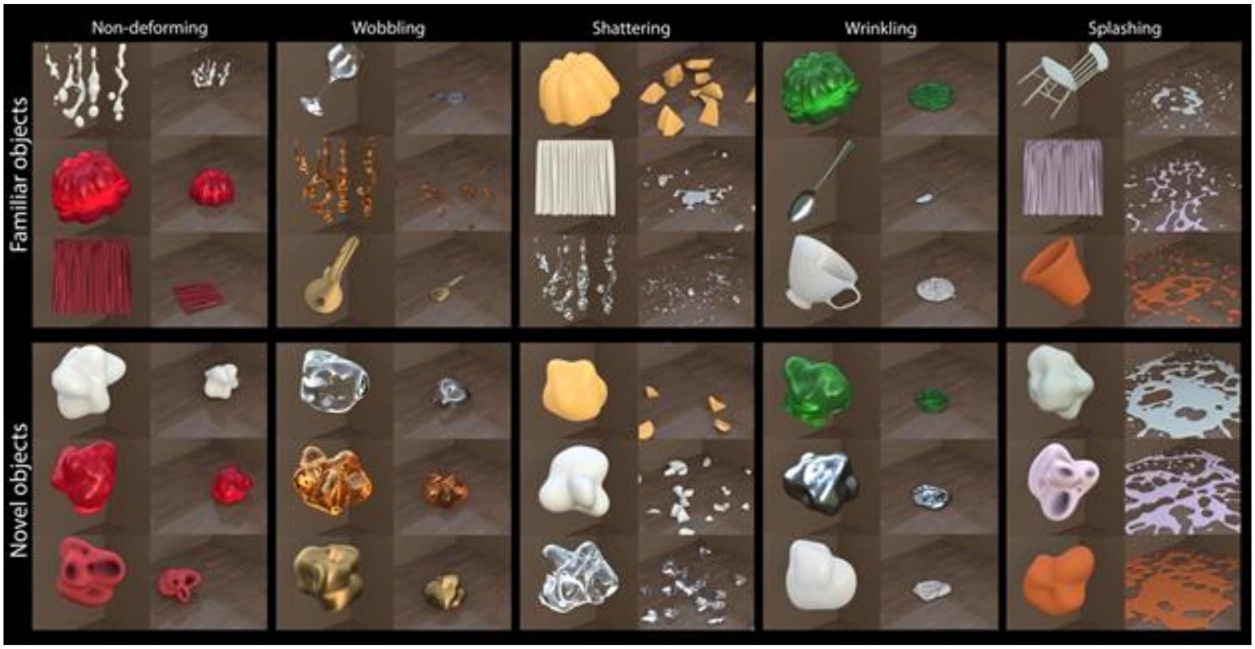


**Supplementary Figure 1. Stimuli Surprising Motion.** Shown are familiar (top) and corresponding novel objects (bottom) used in the Surprising condition. Here, objects are organized according to their material kinematics, however, now the deformations are atypical, e.g. milk shatters, the chair splashes etc. Note that individual scenes are scaled to maximize the view of the object (First frame, left images in each column), or in order to give a good impression of the material kinematics (Last frame, right images in each column).

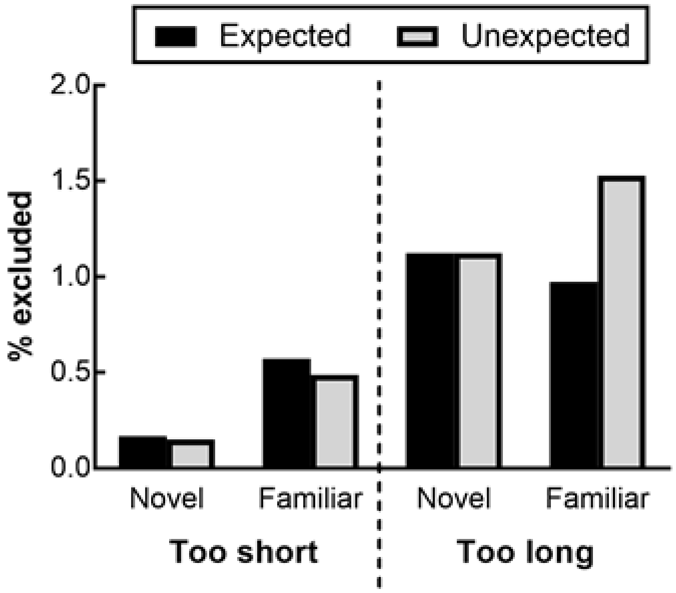


**Supplementary Figure 2**. **RT exclusions.** Trials where Familiar objects that behave unexpectedly are excluded more than other types of trials. It is also interesting that more Familiar objects were excluded for too short RTs – suggests that observers anticipated the rating during the 3 second static hold. Overall only 6% of the data were excluded for reaction times that were too short or too long (~1.3% too fast, ~4.7% too slow. See main text for details.

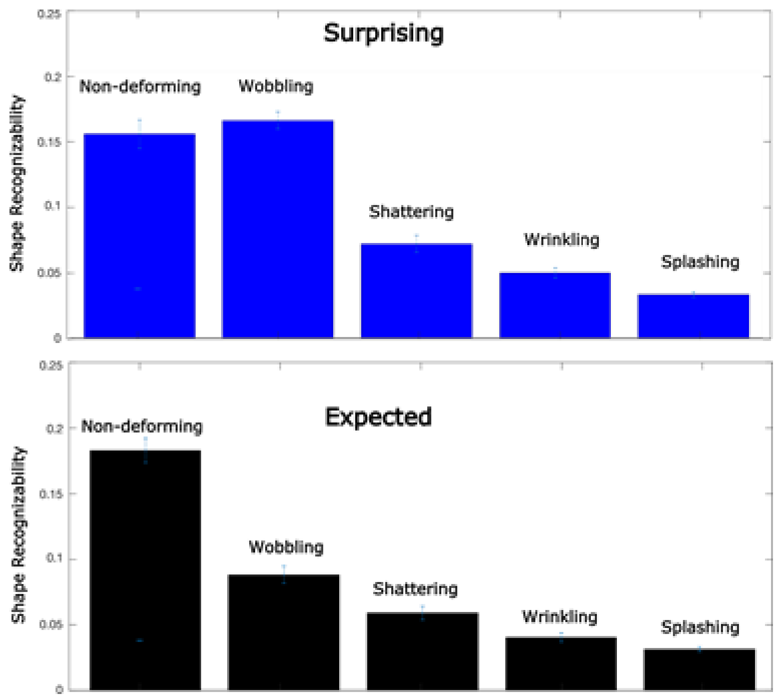


**Supplementary Figure 3. Shape recognizability as a function of material 'behavior' in Surprising and Expected motion condition.** Shape recognizability is defined as the difference between the last frame ratings of familiar and novel objects (weighted by a G-factor as in the regression model in the main text). We believe that the differences in shape recognizability are a direct consequence of how much the object remains intact (unbroken) after hitting the ground. This analysis quantitatively supports the ordering of motions in Figure 7 of the main text. Error bars are 1 SE of the mean.

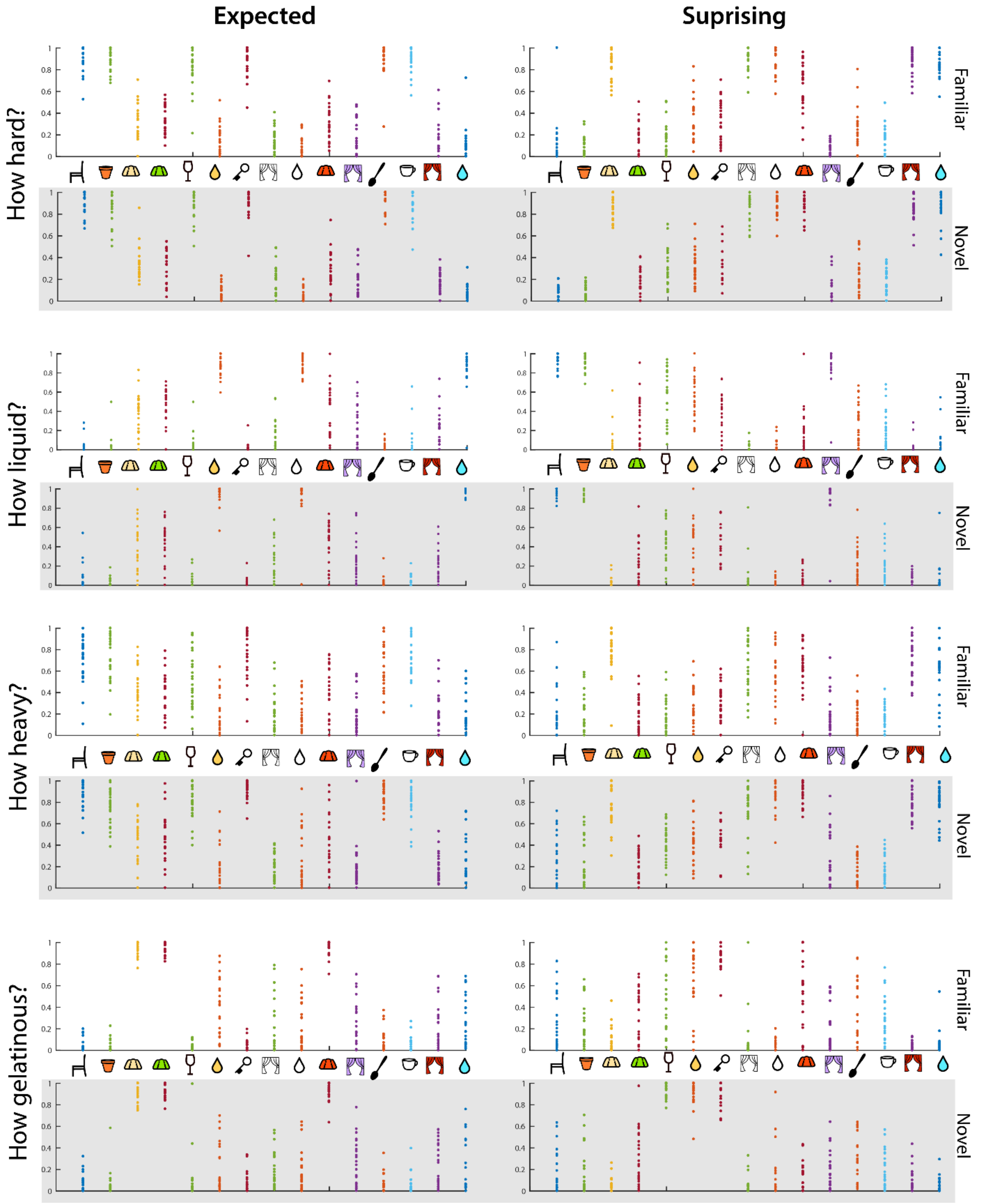


**Supplementary Figure 4. Overall rating distributions.** Shown all observers ratings for each of the 15 familiar objects and corresponding novel objects, all attributes (hard, liquid, heavy, gelatinous) in Expected (left) and Surprising motion conditions (right). It is quite apparent that mean ratings of approximately 0.5 in any condition are not due to 'bimodal' rating behavior, i.e. some observers giving very low and other observers giving very high ratings. Overall observers tended to agree well in their ratings of these four attributes.

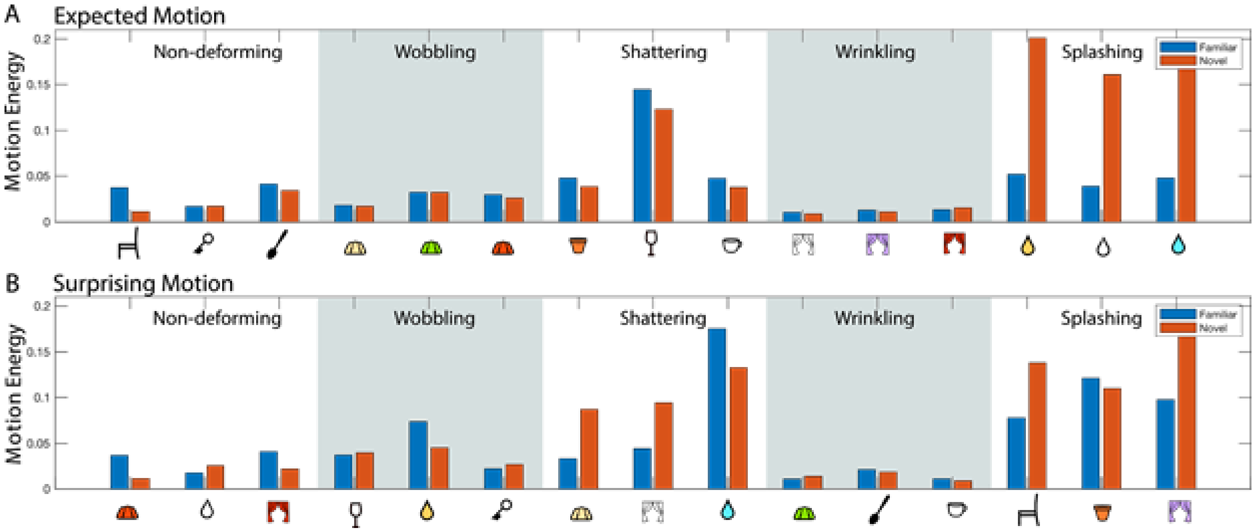


**Supplementary Figure 5. Motion energy for familiar (blue bars) and novel object (orange bars) animations**. Motion energy was computed as the sum of the absolute value of all consecutive image differences, starting with the impact frame (Tschacher, Rees, & Ramseyer, 2014), normalized by the respective. Panel **A** plots the motion energy for familiar and corresponding novel objects in the expected condition. Three objects belong to one motion category (non-deforming, wobbling etc). Similarly, panel **B** plots the motion energy for familiar and corresponding novel objects in the Surprising condition. In the low-level regression model we use the difference in motion energy between familiar and corresponding novel objects as one potential predictor for the observed rating differences between familiar and novel objects.

| | | | \|FF familiar objects - motion novel objects\| | | | | | | Prior Pull= \|Moving Familiar - Corresp. Novel Objects\| | | | | | |
|---|---|---|---|---|---|---|---|---|---|---|---|---|---|---|
| | | | EXPECTED | | | SURPRISING | | | EXPECTED | | | SURPRISING | | |
| DEFORMATION | BLOCK | | Sig. diff | p-value | alpha | Sig. diff | p-value | alpha | Sig. diff | p-value | alpha | Sig. diff | p-value | alpha |
| rigid | hard | milk | **0,072** | **0,000** | 0,0095 | **0,762** | **0,000** | 0,0095 | **0,041** | **0,015** | 0,0280 | 0,004 | 0,794 | 0,0280 |
| | | red Jello | **0,182** | **0,000** | 0,0095 | **0,782** | **0,000** | 0,0095 | 0,017 | 0,673 | 0,0280 | **0,247** | **0,000** | 0,0280 |
| | | velvet | 0,026 | 0,215 | 0,0095 | **0,643** | **0,000** | 0,0095 | 0,011 | 0,430 | 0,0280 | 0,055 | 0,168 | 0,0280 |
| | gelatinous | milk | 0,087 | 0,041 | 0,0095 | **0,150** | **0,003** | 0,0095 | **0,062** | **0,008** | 0,0280 | 0,086 | 0,069 | 0,0280 |
| | | red Jello | 0,033 | 0,071 | 0,0095 | **0,779** | **0,000** | 0,0095 | 0,020 | 0,196 | 0,0280 | **0,392** | **0,000** | 0,0280 |
| | | velvet | **0,202** | **0,000** | 0,0095 | 0,044 | 0,067 | 0,0095 | 0,008 | 0,881 | 0,0280 | 0,055 | 0,036 | 0,0280 |
| | heavy | milk | 0,154 | 0,010 | 0,0095 | **0,754** | **0,000** | 0,0095 | 0,041 | 0,289 | 0,0280 | **0,274** | **0,000** | 0,0280 |
| | | red Jello | 0,098 | 0,076 | 0,0095 | **0,522** | **0,000** | 0,0095 | 0,027 | 0,760 | 0,0280 | **0,238** | **0,000** | 0,0280 |
| | | velvet | **0,095** | **0,001** | 0,0095 | **0,511** | **0,000** | 0,0095 | 0,031 | 0,689 | 0,0280 | **0,088** | **0,003** | 0,0280 |
| | liquid | milk | 0,020 | 0,625 | 0,0095 | **0,881** | **0,000** | 0,0095 | 0,034 | 0,519 | 0,0280 | 0,009 | 0,344 | 0,0280 |
| | | red Jello | 0,032 | 0,449 | 0,0095 | **0,339** | **0,000** | 0,0095 | 0,014 | 0,782 | 0,0280 | **0,130** | **0,009** | 0,0280 |
| | | velvet | **0,096** | **0,009** | 0,0095 | **0,046** | **0,001** | 0,0095 | 0,001 | 0,979 | 0,0280 | 0,014 | 0,187 | 0,0280 |
| wobble | hard | glass | **0,062** | **0,046** | 0,0464 | **0,499** | **0,000** | 0,0464 | 0,032 | 0,810 | 0,0037 | 0,133 | 0,020 | 0,0037 |
| | | honey | **0,058** | **0,000** | 0,0464 | **0,182** | **0,000** | 0,0464 | 0,070 | 0,045 | 0,0037 | 0,040 | 0,263 | 0,0037 |
| | | key | 0,025 | 0,386 | 0,0464 | **0,569** | **0,000** | 0,0464 | 0,002 | 0,698 | 0,0037 | 0,015 | 0,255 | 0,0037 |
| | gelatinous | glass | 0,012 | 0,784 | 0,0464 | **0,884** | **0,000** | 0,0464 | 0,051 | 0,208 | 0,0037 | **0,447** | **0,000** | 0,0037 |
| | | honey | **0,232** | **0,000** | 0,0464 | **0,501** | **0,000** | 0,0464 | **0,167** | **0,001** | 0,0037 | **0,301** | **0,001** | 0,0037 |
| | | key | 0,010 | 0,673 | 0,0464 | **0,801** | **0,000** | 0,0464 | 0,041 | 0,093 | 0,0037 | 0,017 | 0,386 | 0,0037 |
| | heavy | glass | **0,443** | **0,000** | 0,0464 | 0,072 | 0,037 | 0,0464 | **0,282** | **0,000** | 0,0037 | 0,230 | 0,000 | 0,0037 |
| | | honey | 0,057 | 0,191 | 0,0464 | **0,281** | **0,000** | 0,0464 | 0,006 | 0,559 | 0,0037 | **0,124** | **0,004** | 0,0037 |
| | | key | **0,518** | **0,000** | 0,0464 | 0,018 | 0,651 | 0,0464 | 0,195 | 0,004 | 0,0037 | 0,104 | 0,141 | 0,0037 |
| | liquid | glass | 0,008 | 0,566 | 0,0464 | **0,404** | **0,000** | 0,0464 | 0,016 | 0,258 | 0,0037 | 0,123 | 0,067 | 0,0037 |
| | | honey | 0,028 | 0,157 | 0,0464 | **0,478** | **0,000** | 0,0464 | 0,065 | 0,050 | 0,0037 | 0,103 | 0,012 | 0,0037 |
| | | key | 0,008 | 0,407 | 0,0464 | **0,418** | **0,000** | 0,0464 | 0,002 | 0,859 | 0,0037 | 0,039 | 0,579 | 0,0037 |
| shatter | hard | custard | **0,181** | **0,000** | 0,0019 | **0,701** | **0,000** | 0,0019 | 0,021 | 0,841 | 0,0037 | 0,034 | 0,305 | 0,0037 |
| | | linen | 0,081 | 0,009 | 0,0019 | 0,755 | 0,000 | 0,0019 | 0,051 | 0,109 | 0,0037 | 0,024 | 0,343 | 0,0037 |
| | | water | **0,088** | **0,000** | 0,0019 | 0,724 | 0,000 | 0,0019 | 0,042 | 0,223 | 0,0037 | 0,011 | 0,278 | 0,0037 |
| | gelatinous | custard | **0,158** | **0,000** | 0,0019 | 0,719 | 0,000 | 0,0019 | 0,030 | 0,008 | 0,0037 | 0,031 | 0,072 | 0,0037 |
| | | linen | **0,146** | **0,000** | 0,0019 | 0,009 | 0,621 | 0,0019 | 0,043 | 0,349 | 0,0037 | 0,039 | 0,336 | 0,0037 |
| | | water | 0,027 | 0,587 | 0,0019 | **0,177** | **0,000** | 0,0019 | 0,090 | 0,032 | 0,0037 | 0,003 | 0,792 | 0,0037 |
| | heavy | custard | 0,115 | 0,011 | 0,0019 | **0,403** | **0,000** | 0,0019 | 0,030 | 0,356 | 0,0037 | 0,031 | 0,217 | 0,0037 |
| | | linen | 0,041 | 0,137 | 0,0019 | **0,511** | **0,000** | 0,0019 | 0,029 | 0,194 | 0,0037 | 0,101 | 0,050 | 0,0037 |
| | | water | 0,085 | 0,090 | 0,0019 | **0,617** | **0,000** | 0,0019 | 0,037 | 0,086 | 0,0037 | **0,187** | **0,004** | 0,0037 |
| | liquid | custard | 0,055 | 0,310 | 0,0019 | **0,325** | **0,000** | 0,0019 | 0,036 | 0,391 | 0,0037 | 0,032 | 0,161 | 0,0037 |
| | | linen | **0,138** | **0,002** | 0,0019 | 0,014 | 0,691 | 0,0019 | 0,064 | 0,064 | 0,0037 | 0,039 | 0,268 | 0,0037 |
| | | water | **0,029** | **0,001** | 0,0019 | **0,895** | **0,000** | 0,0019 | 0,116 | 0,008 | 0,0037 | 0,007 | 0,750 | 0,0037 |
| wrinkle | hard | green Jello | **0,145** | **0,000** | 0,0016 | 0,004 | 0,885 | 0,0016 | 0,009 | 0,660 | 0,0274 | 0,007 | 0,418 | 0,0274 |
| | | spoon | 0,047 | 0,029 | 0,0016 | **0,651** | **0,000** | 0,0016 | 0,040 | 0,371 | 0,0274 | 0,042 | 0,609 | 0,0274 |
| | | teacup | 0,065 | 0,034 | 0,0016 | **0,653** | **0,000** | 0,0016 | 0,012 | 0,791 | 0,0274 | 0,002 | 0,722 | 0,0274 |
| | gelatinous | green Jello | **0,049** | **0,002** | 0,0016 | **0,584** | **0,000** | 0,0016 | 0,021 | 0,031 | 0,0274 | 0,039 | 0,964 | 0,0274 |
| | | spoon | 0,008 | 0,642 | 0,0116 | **0,219** | **0,000** | 0,0116 | 0,019 | 0,417 | 0,0274 | 0,094 | 0,275 | 0,0274 |
| | | teacup | 0,026 | 0,180 | 0,0016 | **0,116** | **0,002** | 0,0016 | 0,013 | 0,396 | 0,0274 | 0,101 | 0,001 | 0,0274 |
| | heavy | green Jello | 0,167 | 0,003 | 0,0016 | **0,091** | **0,003** | 0,0016 | 0,073 | 0,221 | 0,0274 | 0,028 | 0,378 | 0,0274 |
| | | spoon | **0,504** | **0,000** | 0,0016 | **0,199** | **0,000** | 0,0016 | **0,266** | **0,000** | 0,0274 | 0,006 | 0,752 | 0,0274 |
| | | teacup | **0,339** | **0,000** | 0,0016 | **0,281** | **0,000** | 0,0016 | 0,073 | 0,041 | 0,0274 | 0,006 | 0,389 | 0,0274 |
| | liquid | green Jello | 0,059 | 0,207 | 0,0016 | **0,257** | **0,000** | 0,0016 | 0,022 | 0,246 | 0,0274 | 0,063 | 0,132 | 0,0274 |
| | | spoon | 0,010 | 0,387 | 0,0016 | **0,159** | **0,000** | 0,0016 | 0,001 | 0,910 | 0,0274 | 0,095 | 0,027 | 0,0274 |
| | | teacup | 0,004 | 0,748 | 0,0016 | **0,171** | **0,000** | 0,0016 | 0,034 | 0,260 | 0,0274 | 0,020 | 0,659 | 0,0274 |
| melt | hard | chair | 0,085 | 0,003 | 0,0016 | **0,743** | **0,000** | 0,0016 | 0,008 | 0,367 | 0,0027 | 0,061 | 0,155 | 0,0027 |
| | | clay pot | 0,029 | 0,359 | 0,0016 | **0,827** | **0,000** | 0,0016 | 0,044 | 0,390 | 0,0027 | 0,039 | 0,109 | 0,0027 |
| | | silk | 0,090 | 0,009 | 0,0016 | 0,015 | 0,625 | 0,0016 | 0,026 | 0,231 | 0,0027 | 0,035 | 0,269 | 0,0027 |
| | gelatinous | chair | 0,003 | 0,871 | 0,0016 | 0,090 | 0,032 | 0,0016 | 0,031 | 0,090 | 0,0027 | 0,056 | 0,061 | 0,0027 |
| | | clay pot | 0,017 | 0,507 | 0,0016 | 0,109 | 0,023 | 0,0016 | 0,019 | 0,426 | 0,0027 | 0,068 | 0,049 | 0,0027 |
| | | silk | **0,171** | **0,001** | 0,0016 | **0,104** | **0,009** | 0,0016 | 0,011 | 0,889 | 0,0027 | 0,007 | 0,521 | 0,0027 |
| | heavy | chair | **0,107** | **0,001** | 0,0016 | **0,502** | **0,000** | 0,0016 | 0,137 | 0,003 | 0,0027 | 0,051 | 0,404 | 0,0027 |
| | | clay pot | **0,121** | **0,001** | 0,0016 | **0,401** | **0,000** | 0,0016 | 0,006 | 0,881 | 0,0027 | 0,002 | 0,910 | 0,0027 |
| | | silk | 0,022 | 0,593 | 0,0016 | 0,100 | 0,075 | 0,0016 | 0,007 | 0,877 | 0,0027 | 0,063 | 0,104 | 0,0027 |
| | liquid | chair | 0,007 | 0,803 | 0,0016 | **0,903** | **0,000** | 0,0016 | 0,032 | 0,195 | 0,0027 | 0,029 | 0,095 | 0,0027 |
| | | clay pot | 0,017 | 0,074 | 0,0016 | **0,958** | **0,000** | 0,0016 | 0,003 | 0,029 | 0,0027 | 0,038 | 0,007 | 0,0027 |
| | | silk | **0,149** | **0,002** | 0,0016 | **0,848** | **0,000** | 0,0016 | 0,021 | 0,545 | 0,0027 | 0,025 | 0,663 | 0,0027 |

**Supplementary Table 1. Statistical results for rating differences; FF familiar objects and motion novel objects and prior pull.** This table highlights significant absolute rating differences between **FF familiar objects and motion novel objects (left blue 2 columns) and moving familiar and corresponding novel objects (prior pull, right gray 2 columns),** arranged by deformation method, rating question, and object identity. Values are FDR-corrected, based on a family-wise alpha of .05, for 24 tests (for each deformation: 12 tests for each of the Expected and Surprising conditions). Columns represent differences, p-values, and FDR alpha values for both Expected and Surprising conditions. Bolded values indicate statistically significant differences (also see Figure 7).
